# Histone H2A.X contributes to Gene Expression Regulation and Cell-Cycle Progression in *Toxoplasma gondii* Tachyzoites

**DOI:** 10.64898/2026.10.08.757790

**Authors:** Constanza Cristaldi, Sofia Horjales, Leonardo Gualano, Agustina Ganuza, Ayelén A. Aparicio Arias, Luiz Daniel de Barros, William J. Sullivan, Andrés M. Alonso, Sergio O. Angel, Maria E. Francia, Laura Vanagas

## Abstract

Histone H2A.X is a conserved variant classically associated with the DNA damage response (DDR), where its phosphorylation (γH2A.X) marks double-strand breaks and recruits repair factors. H2A.X is absent in many parasitic protozoans such as *Plasmodium*, *Trypanosoma* and *Leishmania*. In *Toxoplasma gondii*, the obligate intracellular parasite, causative agent of toxoplasmosis, however, its presence raises the question of whether it contributes to genome stability. To tackle this, we generated a *T. gondii* knockout of H2A.X (RHΔH2A.X) and evaluated its phenotype. Although H2A.X is phosphorylated in response to DNA damage in *T. gondii* as in other systems, RHΔH2A.X parasites showed no increased sensitivity to genotoxic agents, indicating that H2A.X is dispensable for canonical DDR. Instead, loss of H2A.X caused reduced gliding motility, increased host cell invasion and clustering of extracellular parasites after egress. Despite no defects in replication rate, RHΔH2A.X showed delayed mitotic progression with abnormal vacuole morphology. Transcriptomic profiling revealed a moderate yet coordinated reprogramming of protein-coding genes. Specifically, H2A.X loss altered the expression of key apical secretory and virulence factors, especially ROP18, while inducing the transcriptional activation of mitotic and DNA replication machinery. Differential enrichment analysis further revealed that upregulated genes were preferentially associated with S phase, whereas downregulated genes were enriched during mitosis and cytokinesis. This is consistent with the delayed mitotic progression and spindle abnormalities observed by microscopy and flow cytometry and suggests a cell cycle phase-specific mechanism of H2A.X-dependent transcriptional control. Rather than a driver of DDR, these findings suggest that *T. gondii* H2A.X operates as a chromatin organizer that coordinates cell division fidelity and virulence gene expression.

**Author Summary:** Cells package DNA into chromatin, composed of histone proteins, whose variants fine-tune gene activity in response to cellular needs. One such variant, H2A.X, is best known across eukaryotes for its canonical role in signaling DNA damage: upon double-strand breaks, H2A.X is phosphorylated and acts as a molecular beacon to recruit repair machinery. Intriguingly, H2A.X is absent in many parasites closely related to *Toxoplasma gondii*, the opportunistic pathogen responsible for toxoplasmosis. We therefore explored the functional role of H2A.X in *Toxoplasma*. By generating parasites completely lacking this histone variant, we found that H2A.X is dispensable for the canonical DNA damage response. Instead, H2A.X operates as a global chromatin organizer that controls the timing of cell-cycle progression and regulates the expression of genes essential for parasite motility, host cell invasion, and virulence. Our findings reveal that *Toxoplasma* has repurposed a conserved histone variant for functions beyond DNA repair, with potential consequences for how this parasite establishes and maintains infection.

## 1. Introduction

*Toxoplasma gondii* is an obligate intracellular apicomplexan parasite and the causative agent of toxoplasmosis. Transmission occurs through ingestion of oocysts in the environment, contaminated food/water, or vertical transmission from mother to fetus [1]. Congenital infection can cause severe neurological damage and chorioretinitis in newborns, with sequelae that may appear years after birth. Infection is often asymptomatic in immunocompetent individuals, but the parasite converts into latent tissue cysts that can reactivate to cause life-threatening damage in immunocompromised patients [2]. Approximately one-third of the global population is infected [3]. The widespread distribution of *T. gondii* relies on its ability to infect virtually all mammals and birds and to alternate between rapidly replicating tachyzoites, slow-growing bradyzoites that form tissue cysts, and environmentally stable oocysts produced through sexual reproduction in felines. This developmental plasticity depends on precise regulation of gene expression [4], primarily orchestrated by ApiAP2 transcription factors together with a Myb-like family transcription factor BFD1 and RNA-binding protein BFD2/ROCY1 [5–7].

Chromatin structure also plays a central role in regulating gene expression. Histones are the main structural components of chromatin, highly conserved proteins responsible for packaging DNA within the nucleus of eukaryotic cells [8]. They are the fundamental substrate upon which chromatin-modifying enzymes, remodelers, and reader proteins exert their regulatory functions. Histones are classified into two main types: canonical histones and histone variants. The latter can confer specific structural and functional properties to chromatin, participating in processes such as transcriptional regulation, cell cycle control, chromosome segregation, and the DNA damage response [9, 10].

Histone variant H2A.X carries a conserved SQE motif at the C-terminal end that is phosphorylated at serine 139 (generating γH2A.X) in response to DNA double-strand breaks (DSBs), or replication stress [11], where it serves as a scaffold for DNA damage response (DDR) factors. Beyond its canonical DDR function, growing evidence from multiple organisms indicates that H2A.X occupies broad genomic regions with functions beyond damage signaling, likely contributing to chromatin organization in a damage-independent manner [11–13]. In particular, H2A.X has been implicated in critical roles in safeguarding mitotic segregation fidelity during cell division [14, 15].

In humans and other mammals, H2A.X represents a minor fraction of total H2A, interspersed with canonical H2A within nucleosomes [16, 17]. In contrast, organisms such as *Saccharomyces cerevisiae*, *Entamoeba histolytica*, *Giardia duodenalis* and *Trichomonas vaginalis* appear to possess only a single H2A-type histone featuring an H2A.X-like C-terminal phosphorylation motif. *Plasmodium* spp. lack a recognizable H2A.X ortholog altogether; sequence analysis indicates that this absence may be longstanding, since the canonical PfH2A retains only a divergent internal SQ motif rather than the SQEF signature characteristic of H2A.X [18]. Nevertheless, this divergent SQ motif in canonical PfH2A is phosphorylated following X-ray-induced DNA damage [19]. This suggests that *Plasmodium* retains a phosphorylation-based DDR signal on its canonical H2A, compensating for the absence of a dedicated H2A.X variant. A similar situation occurs in Trypanosomatids, which also lack a bona fide H2A.X but retain a partially homologous C-terminal motif in their canonical H2A [18], with two phosphorylatable residues (T131 and S133), as demonstrated by tandem mass spectrometry [20, 21].

*Toxoplasma gondii* encodes both a canonical H2A (H2A.1) and H2A.X [22]. TgH2A.X retains the ability to be phosphorylated in response to DNA damage in this parasite [22–24], and the core machinery for DSB repair is present[25, 26]. Furthermore, ChIP-seq data indicate that TgH2A.X is enriched at heterochromatic regions, including subtelomeres, centromeres, and large genomic domains, suggesting a broad structural role in chromatin organization beyond damage signaling [27, 28]. Taken together, these observations support the hypothesis that genomic plasticity, specifically regarding structural variation and recombination rates, and the presence of H2A.X may be inversely correlated across parasitic protozoa: organisms with more plastic genomes (*Plasmodium* spp and Trypanosomatids) tend to lack H2A.X, while those with more stable genomes retain it.

Whether H2A.X actively contributes to genomic stability or DNA repair in *T. gondii*, as in model organisms, remains to be established. To directly address this question, we generated an H2A.X knockout strain and characterized its phenotype. Our results show that H2A.X is dispensable for the canonical DNA damage response but causes broad phenotypic alterations, including differences in motility gliding, invasion, parasite cell division and post-egress dispersal. Transcriptional profiling revealed coordinated reprogramming of invasion-, gliding- and virulence-associated genes, as well as mitotic and cell cycle machinery, consistent with the observed phenotypic alterations. These findings reveal a non-canonical role for H2A.X that impacts multiple aspects of the *T. gondii* lytic cycle.

## 2. Results

### 2.1. Generation and validation of the H2A.X knockout strain

To investigate the role of histone H2A.X in *T. gondii*, we generated a knockout strain (RHΔH2A.X) using CRISPR-Cas9 genome editing. Genome-wide CRISPR fitness screens had predicted H2A.X to be dispensable for in vitro tachyzoite growth [25, 29, 30], providing the rationale to pursue a targeted knockout. Hence, to finally investigate the role of histone H2A.X in *T. gondii*, we directed Cas9 to the promoter region of the H2A.X gene, and co-transfected with DHFR pyrimethamine-resistance minigene cassette [31]. This cassette was PCR-amplified, with homology regions at the 5′ end (adjacent to the Cas9 cut site) and at the 3′ end of the H2A.X locus. (Fig. S1A). PCR using genomic DNA and RT-qPCR assays confirmed the loss of both the H2A.X gene and its transcript in the knockout clones (Fig. S1C, D). The absence of H2A.X in selected clones was confirmed by IFA using anti-H2A.X antibody [22] (Fig. 1A). These results were further validated by western blot analysis, showing loss of the ∼17 kDa H2A.X band in the RHΔH2A.X clones (Fig. S1B). These results collectively validate the successful generation of the RHΔH2A.X knockout line and confirm that H2A.X is dispensable for parasite viability under standard in vitro conditions. For further analysis we used one of the clones successfully generated.

**Figure 1.**
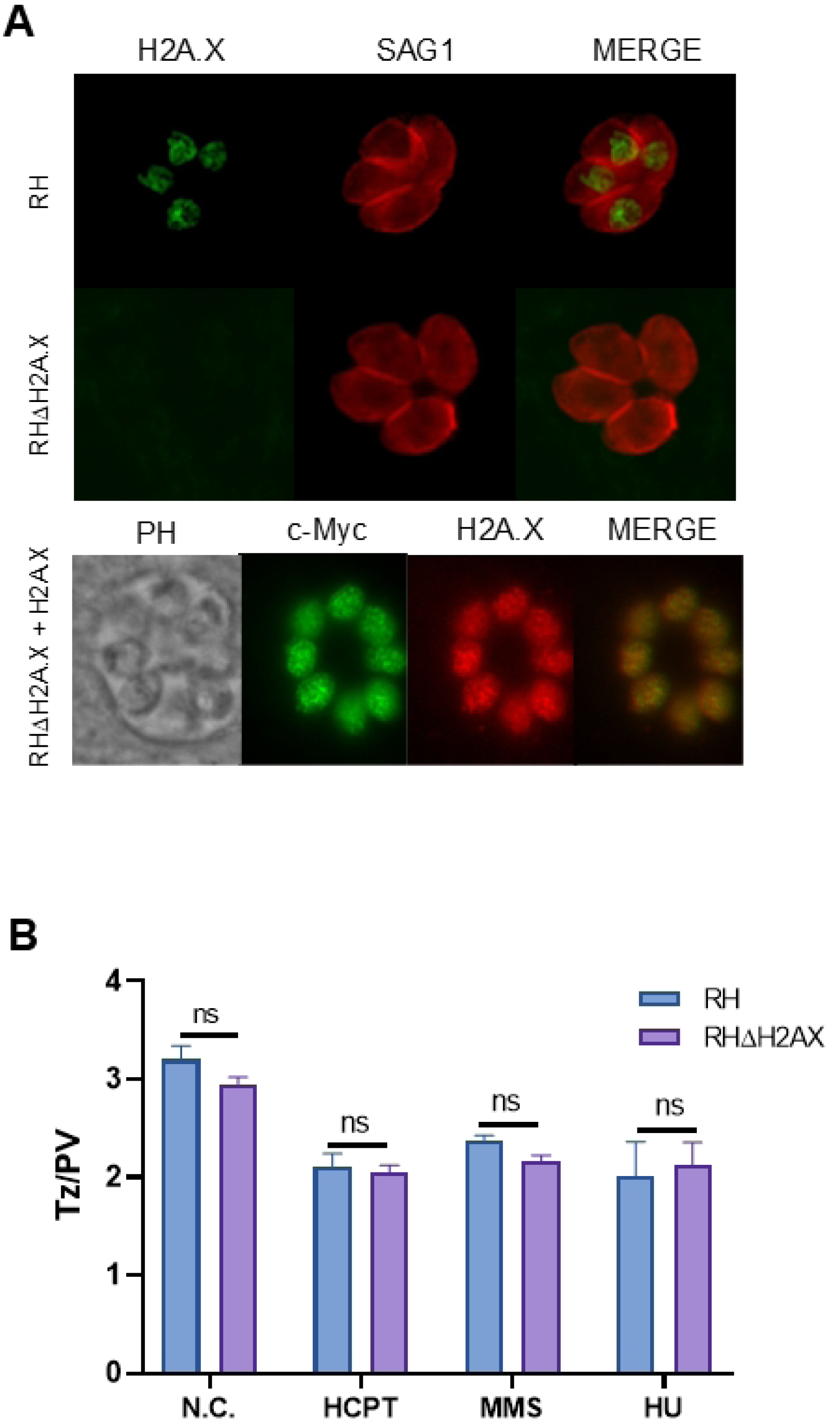
Loss of H2A.X does not sensitize tachyzoites to genotoxic stress. (A) Validation of RHΔH2A.X and RHΔH2A.X+ H2A.X lines by IFA. hTERT cells grown on coverslips were infected with selected clones and the RH parental strain. After 18–24 h, cells were fixed and stained using H2A.X, SAG1 or c-Myc antibodies. H2A.X signal was absent in RHΔH2A.X clones. C-Myc and H2A.X signals co-localized in the nuclei of RHΔH2A.X+ H2A.X tachyzoites. (B) Replication rates were assayed in normal conditions and the presence of drugs that induce DSBs: HCPT (12 μM), MMS (50 μM) and HU (0.2 mM) with no significant differences found between the two strains. N.C.: normal culture conditions. The average number of tachyzoites per PV is graphed.

### 2.2. Sensitivity to genotoxic agents was not altered in RHΔH2A.X tachyzoites

A key mechanism through which this variant histone participates in signaling DNA damage in other species is through phosphorylation of its C-terminal tail serine 139 [16]. γH2A.X mediates recruitment of the repair machinery to the damaged site [32, 33]. This serine has been shown to be phosphorylated in *T. gondii*, and γH2A.X foci increase upon DNA damage [24]. We asked whether parasites lacking H2A.X would display increased sensitivity to genotoxic stress. To address this, we evaluated replication rates after 20 h in both parental and RHΔH2A.X tachyzoites by quantifying the number of tachyzoites per PV under normal conditions (N.C.) and in the presence of different genotoxic drugs. Surprisingly, loss of H2A.X did not affect replication rate either under normal growth conditions or upon genotoxic stress caused by HCPT (Hydroxy-campthotecin, a Topoisomerase I inhibitor) [34], MMS (Methyl-methanosulphonate, which produces DNA damage after methylation) [23] or HU (Hydroxyurea, which produces dNTPs depletion) [35] (Fig. 1B). Consistent with these results, plasmidic DNA integration, which requires activation of DNA repair machinery, was as efficient in RHΔH2A.X as in RH tachyzoites, and stable clones expressing c-Myc tagged H2A.X were successfully generated (see Materials and Methods/Fig. 1A). Stable populations were similarly obtained with three independent plasmids encoding tagged histones (not shown), indicating that loss of H2A.X does not impair DNA-damage facilitated plasmid integration by NHEJ.

### 2.3. Phenotypic alterations after H2A.X loss

Having established that H2A.X is not required for the response to genotoxic stress, we next examined whether its loss would affect other aspects of parasite biology by analyzing key steps of the *T. gondii* lytic cycle, which initiates with host cell invasion, follows with intracellular replication and resumes with egress from the host cell. Plaque assays, measuring repetitive iterations of this cycle, revealed no differences in plaque size (Fig. 2A, B). However, a significantly higher number of lysis plaques in RHΔH2A.X compared to parental RH, suggested increased invasion efficiency (Fig. 2C). Complementation with recombinant, ectopic c-Myc-tagged H2A.X (Fig. 1A, lower panel) restored plaque number to parental levels (Fig. 2C). We next dissected the contributions of invasion, egress, and gliding motility to the observed plaque phenotype.

**Figure 2.**
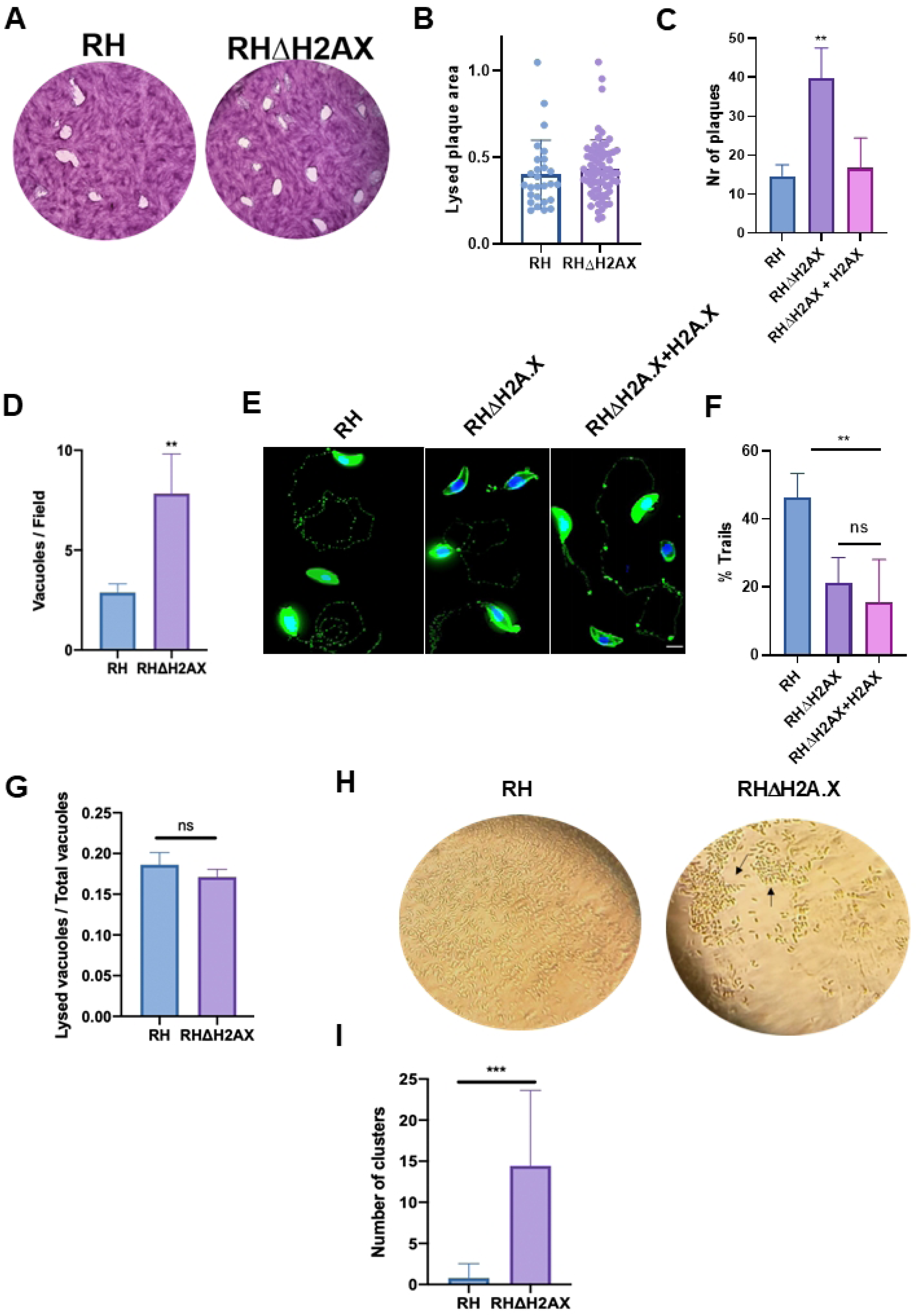
H2A.X loss differentially affects invasion, gliding, and tachyzoite dispersal without impairing egress. (A) The lytic cycle was analyzed after 7 days to allow plaque formation with the RH and RHΔH2A.X strains in confluent hTERT monolayers. Monolayers were fixed and stained with crystal violet. Representative images are shown. (B) Quantitative analysis of plaque area size revealed no significant differences between both strains; however (C) the number of plaques was increased in RHΔH2AX (*Student’s t test,* **p < 0,01). This phenotype was reverted in the complemented line (RHΔH2AX+H2A.X. (D) Invasion rate was evaluated by quantification of the number of vacuoles per field after 30 minutes incubation, exhaustive washing and posterior incubation for 18 h at 37°C (*Student’s t test,* **p < 0,01). (E) Gliding ability was evaluated by IFA using antibodies against SAG1. Representative images are shown for both RH, RHΔH2A.X and complemented (RHΔH2AX+H2A.X) strains. Scale: 5 μm. (F) The number of motile parasites was quantified by Image J. The gliding phenotype of the parental strain was not recovered in the complemented line (*One Way ANOVA, Tukeýs multiple comparison test*, **p < 0,01). (G) Egress was evaluated after incubation with the calcium ionophore (A23187, 2 µM) incubation. The number of lysed vacuoles relative to the total was quantified by IFA, showing no statistical difference. (H) RHΔH2AX parasites showed defective lysis, with the formation of parasite clusters not observed in the parental RH strain. Representative images obtained by photography of recently lysed monolayers in the inverted light microscope are shown. Arrows show characteristic clusters observed after RHΔH2A.X lysis. (I) Clusters were quantified by IFA of extracellular tachyzoites and found to be significantly more frequent in RHΔH2A.X (*Student’s t test* ***p < 0.001). In all cases one representative of three independent assays performed in triplicate is shown.

Invasion was tested by counting PVs after letting tachyzoites invade for 30 min, exhaustive washing, and afterwards over-night replication. RHΔH2A.X parasites exhibited a significantly higher number of PVs than the parental strain (Fig. 2D, p < 0.01), suggesting that H2A.X may be directly or indirectly involved in the invasion process. Because invasion and gliding are closely linked processes in *T. gondii [36]*, we assessed gliding motility assay. RHΔH2A.X parasites exhibited a reduced proportion of motile cells compared to the RH strain. The complemented strain did not fully rescue gliding motility (Fig. 2E, F, p < 0.01).

### 2.4. Tachyzoite lacking H2A.X form aggregates following egress

To assess tachyzoite egress from host cells, intracellular parasites were treated with a calcium ionophore, which triggers motility activation and activates egress. While loss of H2A.X did not affect tachyzoite egress from host cells (Fig. 2G), RHΔH2A.X formed abnormal clusters rather than dispersing as individual zoites upon natural, i.e. not forced by ionophore, egress (Fig. 2H). This was confirmed by immunofluorescence assays of extracellular parasites and quantified compared to the parental (Fig. 2I, p < 0.001). We hereon refer to this phenotype as “clustering”.

### 2.5. H2A.X is required for faithful cell division and mitotic progression

While analyzing the number of tachyzoites per PV, we observed several alterations in parasite morphology, including abnormal arrangement of parasites within vacuoles with loss of the characteristic rosette arrangement, disorganized parasite orientation, frequently uneven parasite numbers per vacuole, and a subpopulation of enlarged binucleated parasites. Upon quantification, we observed that the frequency of abnormal vacuoles was significantly higher in the KO than in the parental strain (Fig. 3A, B; p < 0.01).

**Figure 3.**
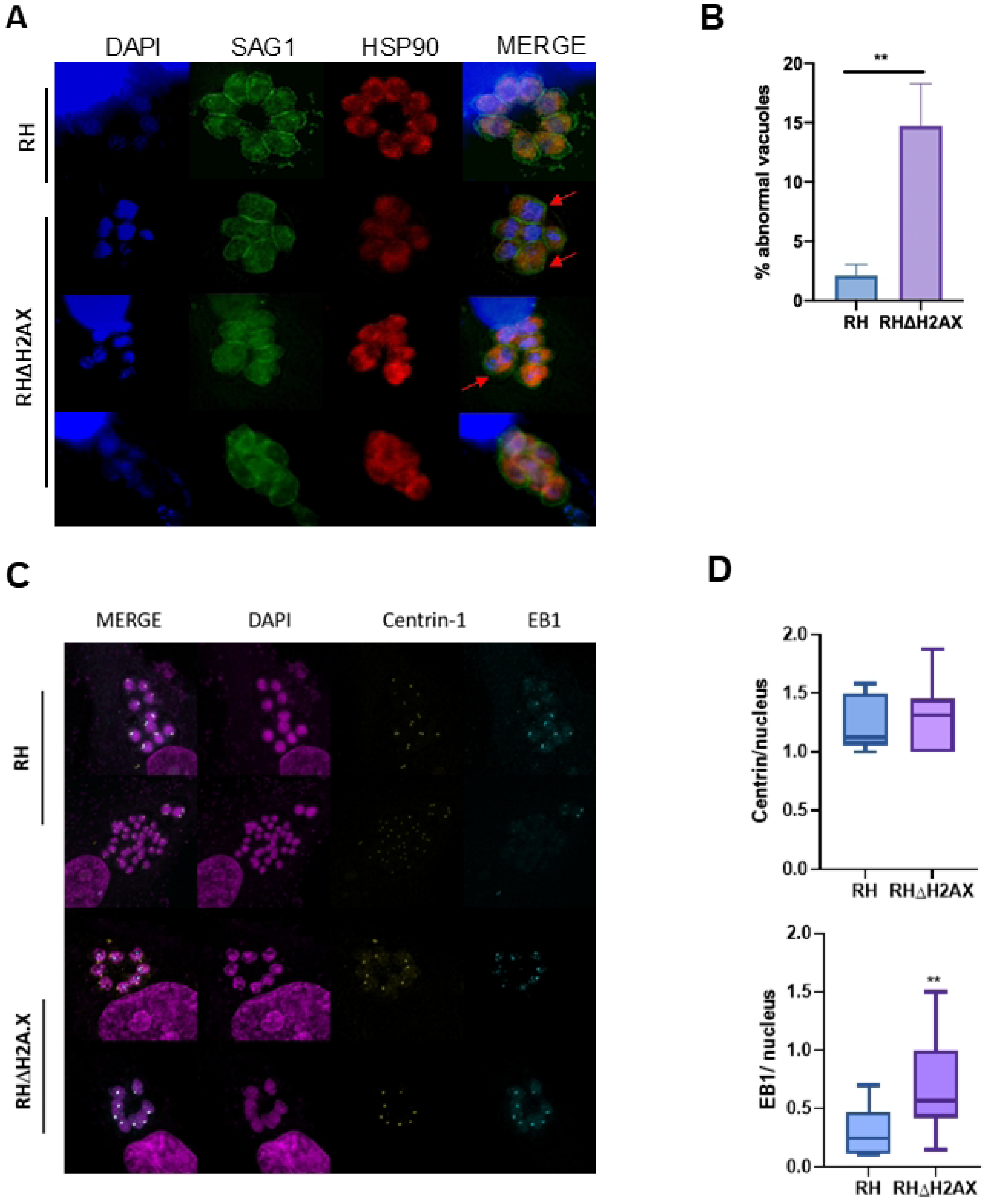
H2A.X histone variant is necessary for normal vacuole morphology in *T. gondii*. (A) IFA using SAG1 (green) and HSP90 (red) antibodies. Abnormal vacuoles were observed in RHΔH2A.X, including enlarged parasites with two nuclei and disorganized vacuoles (arrows). Scale bar: 5 μm. (B) These phenotypes were quantified, showing a significant increase in RHΔH2A.X compared to the parental strain (*Student’s t test* **p < 0.01). (C) IFA performed with EB1 (green) and centrin 1 (yellow) antibodies; representative images. (D) RHΔH2A.X parasites displayed a higher number of EB1 foci than RH and no significant differences in centrin foci (*Student’s t test, \**\*p < 0.01).

To investigate the mechanism underlying this abnormality, we first analyzed the distribution of centrin-1, a bona fide marker of the centrosome [37], EB1, a marker of mitotic spindle [38] and Cpn60, an apicoplast marker [39]. Analysis of centrin-1 and Cpn60 showed no differences in distribution, but a higher proportion of parasites with EB1 foci was observed in RHΔH2A.X (Fig. 3C, D; p<0.01; Fig. S2). The spindle forms upon entry into mitosis, shortly after centrosome duplication, and concomitantly with S-phase progression [40]. However, expansion microscopy revealed no overt differences in subpellicular microtubule organization or mitotic spindle structure (Fig. 4A), indicating that the mitotic defect is not caused by gross cytoskeletal disorganization.

**Figure 4.**
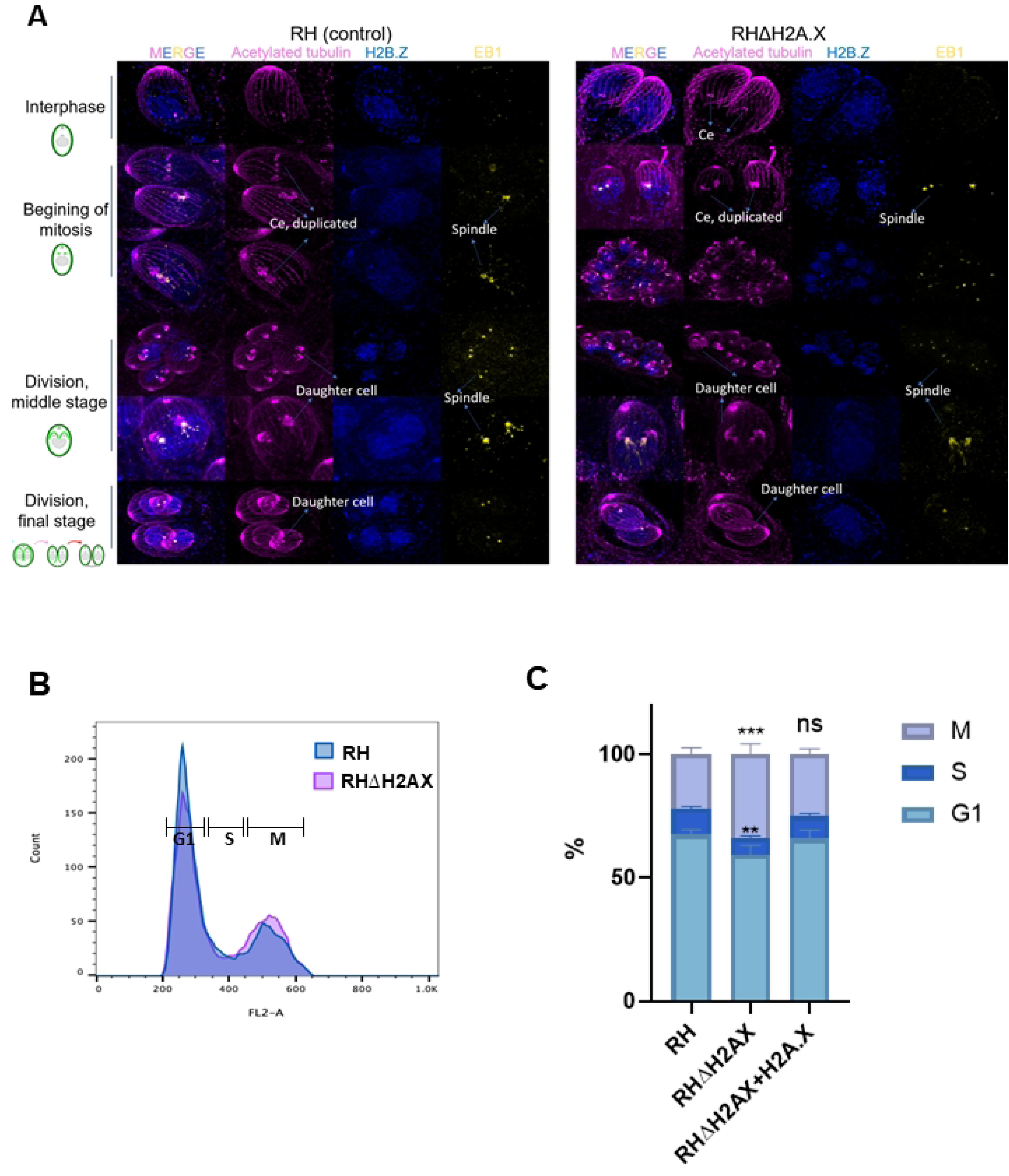
Cytoskeletal disorganization is not responsible for the mitotic delay in the absence of H2A.X. (A) Ultrastructure Expansion Microscopy (U-ExM) was employed to detect potential architectural defects during cycle progression. Representative images of interphase, early, middle and late division stages are shown for both RH (left panels) and RHΔH2A.X (right panels) tachyzoites. Samples were stained for acetylated tubulin (purple), H2B.Z (blue), and EB1 (yellow). Ce; centrosome (B) Cell cycle analysis by flow cytometry, representative histogram showing RHΔH2A.X and RH after propidium iodide staining. Peaks corresponding to G1 – S – M phases are shown and were quantified in (C). RHΔH2A.X + H2A.X complemented strain show no difference compared to the RH strain, while an increased proportion of RHΔH2A.X parasites were found in mitotic (M) phase and concomitantly less proportion of parasites were found in G1 compared to the RH strain (*Student’s t test,* **p < 0,01,***p < 0,001, ns: non significative). Representative of three independent experiments performed in triplicate.

An increased number of EB1 foci could suggest a delay in mitotic progression, or even mitotic arrest. To dissect this, we evaluated the relative frequencies of parasites in different cell cycle stages. Flow cytometry of propidium iodide-stained parasites confirmed accumulation of RHΔH2A.X parasites in mitotic phase, with a concomitant reduction of parasites in G1 (Fig. 4B, C; p<0.01 and p < 0.01 respectively). Complementation with c-Myc-H2A.X restored the normal proportion of mitotic-phase parasites (Fig. 4C), confirming that the cell cycle phenotype is directly attributable to H2A.X loss.

### 2.6. Transcriptomic analysis reveals a global but moderate reprogramming of the tachyzoite transcriptome in the absence of H2A.X

To investigate the transcriptional consequences of H2A.X loss, we performed RNA-seq on intracellular tachyzoites from parental and RHΔH2A.X strains. A three-method intersection approach (DESeq2, edgeR, limma; see Materials and Methods) identified 52 upregulated and 329 downregulated genes shared across all three methods (|log2FC| > 1, *p-adjusted* < 0.05; colored circles in Fig. 5A; 3372 significant DEGs overall, Table S1). In general, transcriptional changes observed are moderate, consistent with the mild but reproducible phenotypic effects observed in RHΔH2A.X parasites, which complete the lytic cycle albeit with reduced efficiency.

**Figure 5.**
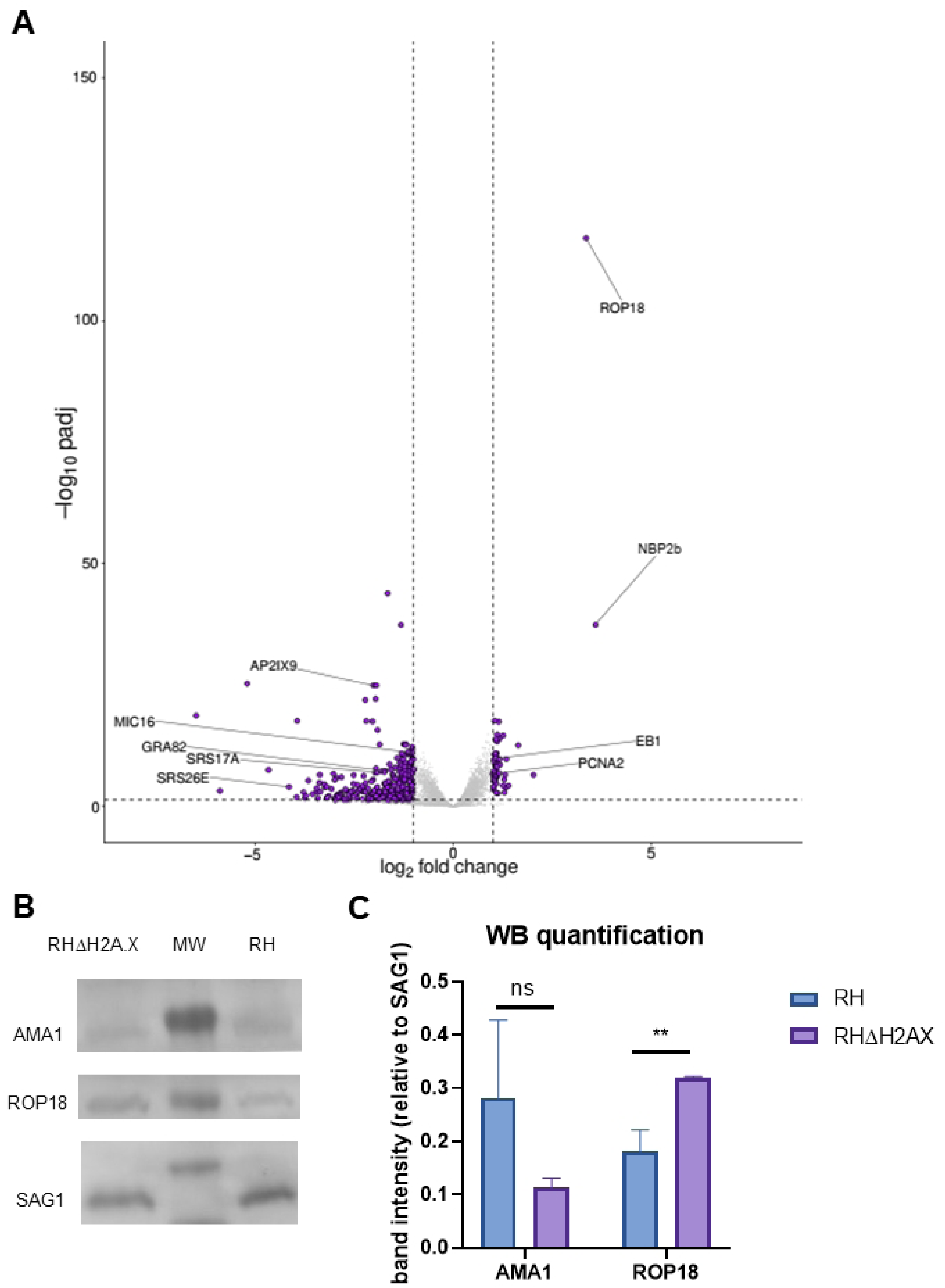
Transcriptomic analysis reveals global reprogramming of tachyzoite identity in RHΔH2A.X tachyzoites. (A) Volcano plot showing differential gene expression in RHΔH2A.X vs. RH. The plot was constructed with significantly enriched genes obtained from DESeq2 analysis. Only genes passing the three-method intersection criterion (padj < 0.05, |LFC| > 1) were highlighted (purple). (B) Western blot analysis of RHΔH2A.X and RH strain abundance of AMA1 and ROP18 proteins. SAG1 antibody was used as charge control and for comparative quantification. MW: molecular weight marker. (C) Quantification was performed with ImageJ. Density of ROP18 and AMA1 bands was relativized to SAG1 on each lane. Three independent experiments were performed, and comparisons by *Student’s t test **p<0.02*.

The most significantly upregulated gene in RHΔH2A.X was the virulence kinase ROP18 (TGME49_205250), confirmed at the protein level by western blot (Fig. 5B, C). Other robustly upregulated genes included an uncharacterized RBL-family protein implicated in host-cell recognition in other apicomplexans (NBP2b-like, TGME49_267875) and the microtubule-associated protein EB1 (TGME49_227650). Because both ROP18 and EB1 show peak expression during S/M phase in cell-cycle transcriptome datasets, their upregulation in bulk RNA-seq must be interpreted alongside the shifted cell-cycle phase distribution of RHΔH2A.X parasites (see Discussion). Among downregulated genes (∼56% hypothetical proteins), some invasion-related genes were prominent (MIC16, GRA82, SRS17A, SRS26E), along with the ApiAP2 factor AP2IX-9 and two hypothetical proteins (TGME49_216335 and TGME49_216290), with predicted membrane-trafficking domains (based on AlphaFold/Foldseek structural predictions) (Fig. 6A; Table S1).

**Figure 6.**
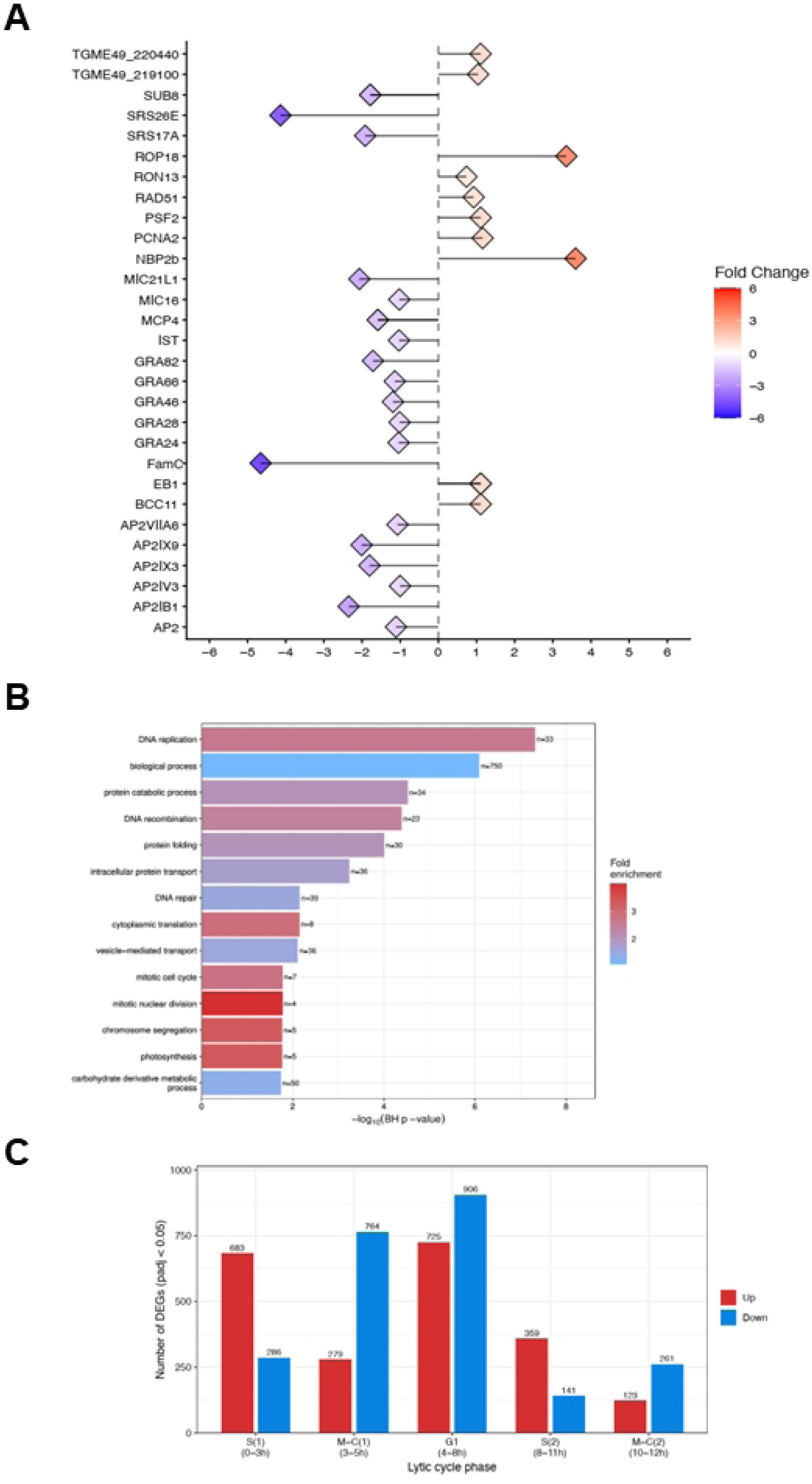
Gene expression profile, biological process enrichment and cell-cycle phase distribution of differentially expressed genes in the RHΔH2A.X mutant. (A) A lollipop plot representing the tendencies of up-regulated and down-regulated genes for RHΔH2A.X. FoldChange: LFC from DESeq2 analysis. (B) Bar plot showing the most enriched biological process for RHΔH2A.X up-regulated genes. Fold enrichment represents the number of times that a term was enriched in our analysis; BH: Bonferroni-Holm correction; n= number of up-regulated genes related to the corresponding biological process. (C) Number of DESeq2 up- (red; log2FC > 0) and down-regulated (blue; log2FC < 0) genes (RHΔH2A.X vs. RH, padj < 0.05) whose hour of peak expression in the reference cell-cycle transcriptome of RH tachyzoites[41] falls within each phase window. Phase windows overlap intentionally at their boundaries, reflecting the gradual nature of lytic-cycle phase transitions.

#### 2.6.1 Coordinated regulation of the invasion and gliding machinery by H2A.X histone

Beyond individual significantly up- or downregulated genes, RHΔH2A.X showed a broader, coordinated remodeling of the secretory/surface protein repertoire: widespread downregulation of microneme, dense granule, and SRS genes, indicating a broad reduction in the expression of parasite surface antigens, along with the already mentioned hypothetical proteins, putatively associated to membrane trafficking (Fig. 6A, Table S2). This coordinated decrease involved multiple functional classes of invasion-associated proteins, including core adhesins, adaptor proteins, subtilisin-like proteases, rhomboid proteases, and the moving junction ligands AMA1 and AMA2. Notably, reduced AMA1 transcript abundance was accompanied by decreased protein expression (Fig. 5B, C). In addition, several calcium-dependent protein kinases that regulate microneme secretion (CDPK2/5/6/8) and the MYR translocon (MYR1–4), accompany this trend (Fig. 6A, Table S2).

In contrast, rhoptry neck genes required for moving-junction assembly (RON2, RON4, RON4L1, RON5, RON8) and rhoptry kinases (including ROP18) were consistently upregulated (Table S2). This reciprocal MIC/GRA/SRS-down vs. RON/ROP-up pattern provides a transcriptional framework for the paradoxical phenotype of RHΔH2A.X: impaired gliding coexisting with enhanced invasion.

#### 2.6.2. Cell-cycle phase-linked transcriptional signatures in RHΔH2A.X

GO enrichment of upregulated genes identified DNA replication (GO:0006260) as the top Biological Process term, reflecting a broad cluster of replication factors (PCNA1/2, GINS subunits Psf2/Psf3, an RFA subunit, both RNR subunits, dUTPase, five MCM helicase subunits, two CDK regulatory subunits, and RAD51) (Fig. 6A–B, Table S3). DNA repair and recombination terms (GO:0006281 and GO:0006310, respectively) were independently enriched among upregulated genes, as were translation/ribosomal biogenesis terms. Chromosome segregation (GO:0006260) and mitotic process terms (GO:0000278, GO:0140014) were also enriched (Fig 6B), driven by kinetochore components, condensin subunits, the chromosomal passenger complex (INCENP1/2), ARK kinases, centrosomal proteins, EB1, and the conoid protein BCC11 (Table S3).

To test whether these changes reflect a defined cell-cycle mechanism, we mapped DESeq2-derived up- and downregulated genes (padj < 0.05) onto the RH cell-cycle transcriptome[41] by phase of peak expression (Fig. 6C). While the largest fraction of both up- (log2FC >0) and downregulated (log2FC < 0) genes fell in G1 (4–8 h), genes normally peaking in S phase (S1 0–3 h, S2 8–11 h) were preferentially upregulated in RHΔH2A.X, whereas genes normally peaking during mitosis/cytokinesis (M/C1 3–5 h, M/C2 10–12 h) were preferentially downregulated. This reciprocal S-up/M-C-down pattern is consistent with the delayed mitotic progression phenotype described in section 2.5 (microscopy, flow cytometry), reflecting either altered phase composition, genuine phase-specific transcriptional dysregulation, or both.

#### 2.6.3. H2A.X occupancy does not predict all transcriptional changes

To test whether transcriptional changes reflect direct H2A.X binding, we intersected RHΔH2A.X RNA-seq results with our previously published H2A.X ChIP-seq data [27]. H2A.X occupancy around transcription start sites (TSS) did not differ between up-, down-, and non-regulated genes (Fig. 7A), nor was it enriched among differentially expressed genes relative to the genome-wide background (Fisher’s exact test, p > 0.1 for both directions: |log2FC| < 0 |log2FC| > 0; p-adj < 0.05; Fig. 7B, Table S1), suggesting that H2A.X-driven transcriptional changes are largely indirect. Consistent with this, upregulated genes were significantly enriched among those carrying active promoter marks (H2B.Z, H2A.Z, H3K4me3) in the parental strain [27], while downregulated genes were depleted of the same marks (Fig. S3). This points to a chromatin-context-dependent effect in which H2A.X loss preferentially derepresses genes already in permissive chromatin environments, rather than a locus-specific regulator.

**Figure 7.**
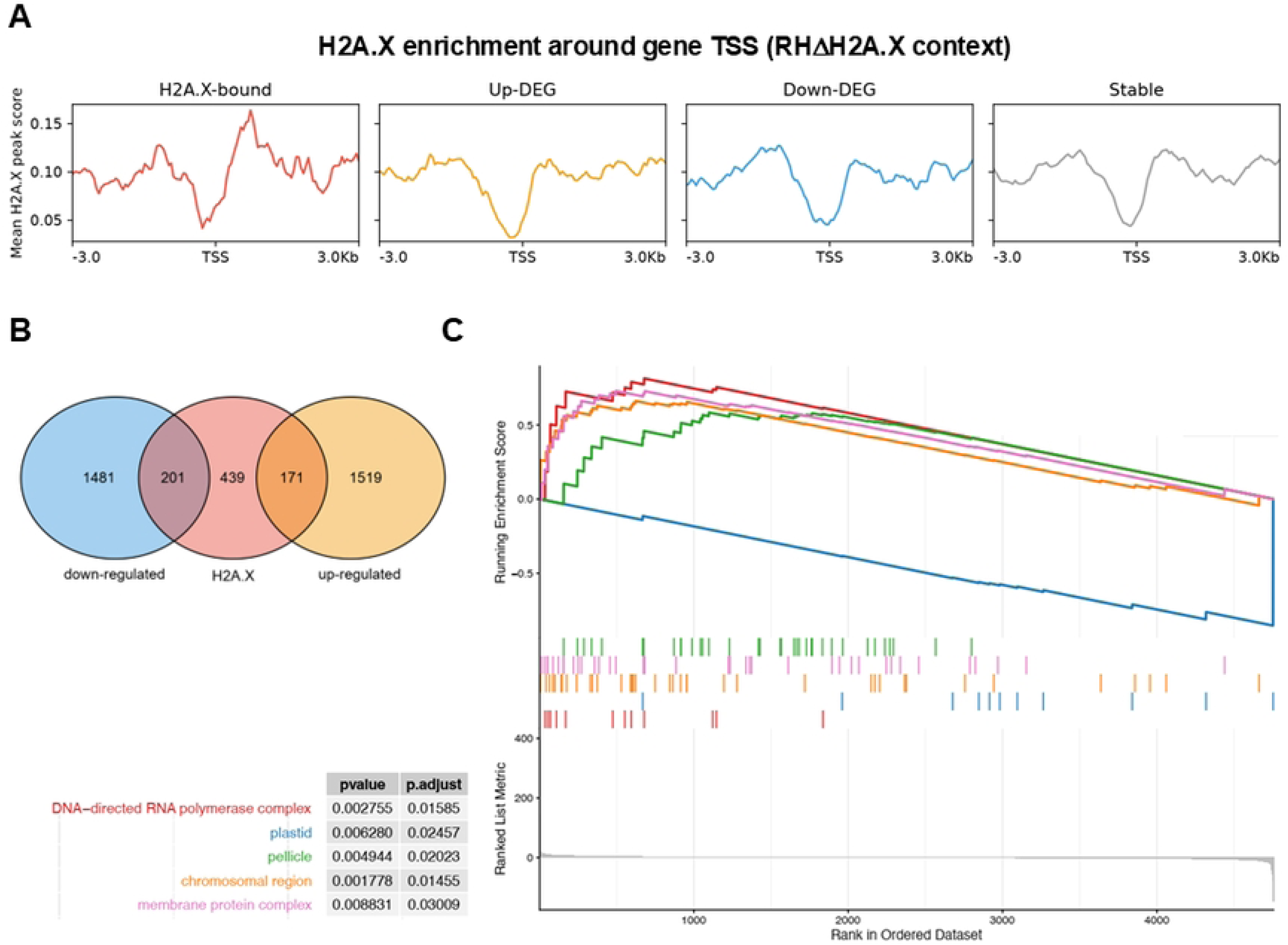
Convergence of H2A.X chromatin occupancy and transcriptional dysregulation in RHΔH2A.X tachyzoites. (A) Metagene H2A.X signal profile around gene TSS, stratified by RHΔH2A.X status. The plot shows the mean H2A.X ChIP signal (peak score) across a ±3 kb window centered on the TSS, computed separately for four gene sets: H2A.X-bound genes, up-regulated DEGs, down-regulated DEGs, and transcriptionally stable genes (reference set of unbound, non-DEG genes). (B) Overlap between H2A.X-bound genes and RHΔH2A.X differentially expressed genes. The three-way Venn diagram partitions the 811 H2A.X-bound genes tested in the DESeq2 analysis against the 1,690 up-regulated and 1,682 down-regulated genes (padj < 0.05, any log2FC). (C) GO Cellular Component GSEA running-enrichment profiles for the five most robust, non-redundant, semantically simplified terms enriched in RHΔH2A.X vs. RH. Top panel: running enrichment score (ES) as a function of gene rank in the ordered list, colored by term -- chromosomal region (orange), DNA-directed RNA polymerase complex (red), pellicle (green), plastid (blue), and membrane protein complex (pink); the inset table reports the nominal p-value and BH-adjusted p-value (padj) for each term. Bottom panel: the ranked-list metric (log2 fold change x -log10(p-value)) for every gene in the pre-ranked list, shown in grey, illustrating the magnitude of differential expression underlying each gene’s rank position.

#### 2.6.4. Subcellular localization of transcriptional changes

Gene set enrichment analysis (GSEA[42]) resulted in only one statistically significant, non-redundant, robust term: Cellular Component (CC) in which five significant, non-redundant terms were included: chromosomal region, DNA-directed RNA polymerase complex, pellicle, plastid/apicoplast, and membrane protein complex (Fig. 7C, Table S1). Four of the five were enriched toward upregulated genes, including a chromosomal-region cluster anchored by the kinetochore component NDC80 (TGME49_260420) and the subtelomeric FamC gene TGME49_200130 (both H2A.X-bound per ChIP-seq), consistent with the delayed mitotic progression phenotype described in section 2.5, and a broad IMC/pellicle biogenesis cluster, consistent with the daughter-cell budding defect described in the same section.

## 3. Discussion

Histone H2A.X is best known as a sensor of DNA double-strand breaks, phosphorylated at its C-terminal SQ motif to form γH2A.X upon genotoxic stress; we showed that this phosphorylation increases in *T. gondii* following DNA damage [24]. Its broad conservation across eukaryotes, however, points to additional, damage-independent functions, including roles in chromosome segregation and the spindle assembly checkpoint (SAC) [43, 44]. Our results are consistent with this view: RHΔH2A.X parasites showed no increased sensitivity to genotoxic agents, indicating that H2A.X is not essential for survival following DNA damage. This does not exclude a role in damage signaling but suggests functional redundancy, potentially with H2A.1 which retains a truncated SQ motif (SQEF), or compensation by parallel repair pathways. Such a supportive, rather than essential, role of H2A.X in DNA damage checkpoint activation and repair, has been proposed previously [45]. Consistent with preserved repair capacity, we generated a stable, drug-resistant complementation line by ectopic reintroduction of c-Myc-tagged H2A.X. Because stable transgene integration in *T. gondii* occurs predominantly through non-homologous integration of the transgene [46], this finding indirectly indicates that NHEJ-mediated genomic integration remains functional in the absence of H2A.X. The evolutionary conservation of H2A.X in *T. gondii* therefore cannot be explained by a role in DNA repair response alone.

The most prominent consequence of H2A.X loss was altered cell-cycle timing: a subpopulation of RHΔH2A.X parasites accumulated in mitosis with abnormal vacuole organization, yet spindle and subpellicular microtubule architecture remained grossly intact. This argues against a primary cytoskeletal defect and instead points to a chromatin-organizer interpretation, which is also supported by GSEA analysis. This analysis revealed coordinated up-regulation of IMC/pellicle biogenesis transcripts, suggesting dysregulation of the daughter-cell budding transcriptional program rather than an intrinsic structural defect. Consistent with this, flow cytometry revealed an increased mitotic subpopulation in RHΔH2A.X parasites, indicative of a partial, non-lethal delay in mitotic progression. Because replication rate was assessed as an endpoint measure (parasites/vacuole at 20 h), this may not translate into a detectable reduction in completed divisions within this window, reconciling the apparent discrepancy between the cytometry and replication-rate data.

Transcriptomic analysis was consistent with the partial mitotic delay observed in RHΔH2A.X. Comparison with the reference tachyzoite cell-cycle transcriptome [41] showed preferential upregulation of genes normally peaking in S phase and downregulation of those peaking during mitosis/cytokinesis. This pattern raises the possibility that H2A.X contributes to the regulation of gene sets associated with specific cell-cycle stages. However, it could also reflect a shift in the phase composition of the asynchronous population rather than genuine transcriptional dysregulation. Individual genes illustrate both possibilities. ROP18 and EB1, which peak during S/M, were upregulated and may therefore reflect the altered cell-cycle distribution. In contrast, AMA1, which peaks in M/C and is lowest in G1/S, was downregulated at both the transcript and protein level, a direction opposite to that predicted by a simple phase-composition shift. Similarly, NBP2b-like TGME49_267875, discussed below, shows no strong cell-cycle-associated expression pattern despite being among the most robustly upregulated genes. This argues that at least part of the transcriptional signature reflects active, H2A.X-dependent regulation rather than population-level redistribution alone, although disentangling the relative contribution of each will require cell-cycle-resolved transcriptomics, an analysis currently underway. A direct regulatory role for H2A.X during mitosis would be consistent with reported functions of γH2A.X in chromosome segregation and kinetochore assembly in other eukaryotes [43, 44], suggesting this connection may be evolutionarily conserved beyond its canonical role in DNA damage signaling.

Transcriptomic and ChIP-seq data together support a model in which H2A.X acts predominantly as a chromatin organizer rather than a locus-specific transcriptional regulator: H2A.X occupancy did not predict whether a gene was up- or downregulated after knockout, yet functionally related gene sets shifted in coordinated fashion. We found a small number of exceptions, including the kinetochore component NDC80 and the subtelomeric FamC member (TGME49_200130), both previously identified as H2A.X ChIP-seq associated peaks in the parental strain. This indicates that direct binding does contribute to a minority of transcriptional changes, while the bulk of the transcriptional response likely arises indirectly. When looking at active promoter marks such as H2B.Z, H2A.Z and H3K4me3 [27], once again the results support a model in which H2A.X acts as a context-dependent modulator of transcription rather than as a global activator or repressor. Although these findings are based on correlative integration of two independent datasets, the consistency across three independent chromatin marks and two distinct genomic compartments (euchromatin and subtelomeric heterochromatin) strengthens the biological relevance of the observed association. ChIP-seq profiling in the knockout background would be required to directly assess chromatin redistribution

Among these phenotypic consequences, H2A.X loss uncouples gliding from invasion, with RHΔH2A.X parasites being less motile yet invading more efficiently. Transcriptional data show downregulation of microneme/motility genes with upregulation of RON genes required for moving-junction assembly. Because RON-dependent tight-junction closure and vacuole sealing can proceed independently of the actomyosin motor [47], enhanced invasion may reflect more efficient moving-junction formation rather than compensation for reduced motility. However, this does not distinguish increased attachment from productive invasion, which we address below. Notably, complementation did not fully rescue gliding motility, potentially reflecting secondary genomic alterations resulting from H2A.X loss that are not reversed by ectopic re-expression, or an effect of the epitope tag, as reported previously [48, 49].

The post-egress clustering phenotype may share a related, membrane-level origin. Although a related post-egress grouping has been recently described in cytokinesis-defective *T. gondii [50]*, RHΔH2A.X parasites complete cytokinesis normally and, unlike those conjoined parasites, invade more efficiently, arguing that the clustering and enhanced invasion in this case are independently driven by transcriptional remodeling. Beyond reduced MIC2/ROM4-dependent adhesin shedding, several strongly regulated genes point to broader membrane remodeling. This includes opposite-direction changes in lipid-transfer proteins (VPS13-domain TGME49_216335/216290, downregulated; START-domain TGME49_239510, upregulated) and in cation/calcium-homeostasis genes. Although individually uncharacterized in *T. gondii*, their coordinated, opposite-direction shifts raise the possibility of an altered plasma membrane composition that favors parasite-parasite adhesion after egress, independently of, or in addition to, reduced adhesin processing. The broad transcriptional downregulation of SRS/SAG proteins in RHΔH2A.X could expose adhesion-competent molecules, providing an additional surface-level basis for post-egress parasite–parasite adhesion. These hypotheses will require direct testing, for instance by lipidomics or membrane-fluidity assays.

Among individual genes, ROP18 stands out both for the magnitude of its change, the most significant upregulated transcript in the dataset, and for its role as a major virulence determinant [51]. This finding must be interpreted alongside the phase-composition caveat discussed above; nonetheless, it gains independent support from our previous work showing that H2B.Z acetylation also regulates ROP18 protein abundance, independently of mRNA level [31]. Together, these findings point to ROP18 as a locus subject to multiple layers of chromatin-based regulation in *T. gondii*, and a useful reporter for future work in this area.

While ROP18 upregulation and the broader transcriptional remodeling described above suggest potential virulence consequences of H2A.X loss, the in vivo relevance remains uncertain. A genome-wide CRISPR screen during acute murine infection placed H2A.X within the near-neutral range of the fitness distribution, though gene-specific data were unavailable for this locus and the reported score reflects the reference-strain default [30]; direct in vivo validation of H2A.X’s contribution to virulence therefore remains an open question.

Loss of H2A.X may also de-repress specific apical organelle genes, a phenomenon linked to invasion gene regulation in other apicomplexans [52]. Among the robustly upregulated genes, an uncharacterized NBP2b-like protein (TGME49_267875) is particularly notable, and as noted above, shows no strong cell-cycle-associated expression pattern in available RNA-seq or scRNA-seq datasets. In *Plasmodium*, NBP/RBP family members localize to the rhoptries and participate in invasion complexes [53]. Given the gliding defect in RHΔH2A.X, its upregulation raises the possibility of a compensatory shift in invasion-related gene expression, though functional characterization of TGME49_267875 will be needed to test this directly.

Overall, our findings indicate that H2A.X contributes to chromatin organization, cell-cycle regulation, and transcriptional control of the lytic cycle, rather than being essential for canonical DNA damage responses. Coordinated changes across functional gene sets may underlie the observed defects in invasion, motility, and egress, reflecting both direct and cell-cycle-dependent effects of H2A.X loss.

## 4. Materials and Methods

### 4.1. Parasite strains and Culture

hTERT (BD Biosciences, San Jose, CA, USA) fibroblast monolayers [54] were infected with RHΔHXGPRT strain parasites [55] and incubated in Dulbecco’s Modified Eagle

Medium (DMEM, GIBCO, ThermoFisher Scientific, Waltham, MA, USA) supplemented with 1% fetal bovine serum, penicillin and streptomycin (100 µg/ml; GIBCO, ThermoFisher Scientific) at 37 °C in a 5% CO₂ atmosphere.

### 4.2. Generation of the RHΔH2A.X strain

A guide RNA (gRNA) targeting the promoter region of the H2AX histone gene was designed using the following oligonucleotides: Fw: 5’-<u>AAGTT</u>GTGTGTCCAGTTCCACGCAAG-3’, Rv: 5’-<u>AAAAC</u>TTGCGTGGAACTGGACACACA-3’. The underlined sequences correspond to BsaI restriction sites. The oligos were annealed and cloned into a plasmid encoding Cas9 previously digested with BsaI, followed by ligation. The correct insertion of the gRNA into the plasmid was verified by PCR and confirmed by sequencing. In parallel, primers were designed to amplify the DHFR resistance gene, incorporating at their 5’ and 3’ ends homology sequences to the H2A.X locus. This design allows for homologous recombination-mediated replacement of the target gene. The primers used were: Fw: 5’-<u>CACAACTGCGAGTTTTCTCGTGTGTACACTCAAATTTCAC</u>AAGCTTCGCCAGGCTGTAAATCC-3’ and Rv: 5’-<u>GTCTTCTTACGACCTCCTGCACCTTTGGCGGACATTTTGA</u>TCATCCTGCAAGTGCATAGAAGGA-3’. The underlined regions correspond to sequences homologous to the H2A.X gene. The resulting amplicon was purified from agarose gel electrophoresis. This purified PCR product, together with the Cas9-gRNA plasmid, were used to transfect RHΔHXGPRT tachyzoites. Once the parasites lysed the host cell monolayer, they were cloned by limiting dilution in media with pyrimethamine (1 µM, Sigma-Aldrich, Saint Louis, MO, USA) to enrich for recombinants that had successfully integrated the replacement cassette. Prior to selection, expression of nuclear-localized Cas9 was confirmed by IFA using an anti-FLAG antibody (Sigma-Aldrich, F1804), which detects the tagged Cas9 protein. The resulting clones were screened by PCR using the following primers designed to distinguish positive from negative clones: H2A.X-upUTR-Fw: 5’-GAACTAAGGGTGAACTGATG-3’, H2A.Xdown-Rv: 5’-GCCAAACACCACGATACTCTT-3’, val-H2A.X-Rv: 5’-CAGGCTCAGAGAAAAGCGTAAGTC-3’.

### 4.3. Complementation of the RHΔH2A.X strain

RH or RHΔH2A.X parasites were transfected with plasmids containing different histone genes fused to different tags, such as c-Myc or Flag, after a tubulin promoter. After transfection, the parasites were used to infect cells grown on slides. 24 hours post-infection, the cells were fixed with 100% cold methanol for 7 minutes at -20 °C. IFA were then performed as described above, using antibodies against the tags: α-c-Myc (rabbit, Abcam ab9106 or mouse, Santa Cruz sc-42) and α-Flag (mouse, Sigma F1804). The number of vacuoles positive for c-Myc or Flag was then quantified, showing no significant differences. In three of the experiments, transfected parasites were selected for three passages with mycophenolic acid (MPA, 25 μg/ml, Sigma-Aldrich) and xanthine (Xan, 50 μg/ml, Sigma-Aldrich) and cloned by limiting dilution. Stable clones were obtained in all cases.

One of the transfections was performed with pTUB8Myc-H2A.X vector: 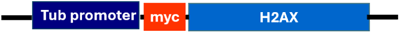 [56], selected for three passages with MPA and Xan and cloned by limiting dilution. Clones of the complemented RHΔH2A.X were obtained (RHΔH2A.X + H2A.X) and checked for phenotypical recovery.

### 4.4. Immunofluorescence assay (IFA)

Intracellular tachyzoites were grown on slides and subsequently fixed with 100% cold methanol for 7 minutes at –20 °C. Extracellular tachyzoites were fixed with PAF 4% in PBS for 30 minutes and placed on slides. After fixation, the samples were washed with PBS and blocked with 3% BSA. IFAs were performed using a mouse anti-SAG1 (1:200, obtained through BEI Resources, NIAID, NIH: Monoclonal Anti-*Toxoplasma gondii* Surface Antigen 1 Protein, Clone T4 1E5 (produced *in vitro*), NR-50255), a rabbit anti-H2A.X (1:200; performed by Eurogentec against unphosphorylated c-terminal TgH2A.X peptide), a rabbit anti-c-Myc (1:200, abCam, ab9106), a rabbit anti-HSP90 (1:500, made in our laboratory), a mouse anti-acetylated tubulin (1:500, Sigma), a rat anti-Ebi1(1:1500, kindly provided by Dr. Marc-Jan Gubbels), a mouse anti-Centrin1 (1:500, Cell Signaling), a rabbit anti-Cpn60 (1:300, kindly provided by Dr. Erica Dos Santos Martins), a rabbit anti-H2B.Z (1:500, made in our laboratory), as primary antibodies for 1 hour at room temperature. After washing, Alexa Fluor-conjugated secondary antibodies were used: goat anti-mouse 594 and goat anti-rabbit 488 or goat anti-mouse 488 and goat anti-rabbit 594 (ThermoFisher Scientific). Nuclei were stained with DAPI. Finally, samples were mounted in Mowiol, and images were acquired using Axio Imager.M2 microscope (Carl Zeiss AG, Oberkochen, Germany) equipped with a Plan-Apochromat 63x/1.40 Oil M27 objective and a 2.8-megapixel monochrome Zeiss 503 digital video camera.

### 4.5. Western blot analysis

Tachyzoites were harvested and lysed by resuspension in Laemmli buffer followed by heating at 95 °C for 10 minutes. Protein samples (5 × 10^6^ to 1 × 10^7^ parasites per well) were separated by 15% or 10% SDS-PAGE and transferred onto a PVDF membrane at 100 V for 1 hour. Non-specific binding sites were blocked with 5% non-fat-dried milk in PBS containing 0.05% Tween-20. Membranes were incubated for 1 hour at room temperature with primary antibodies: rabbit anti-H2A.X (1:5000), mouse anti-SAG1 (1:2000, obtained through BEI Resources, NIAID, NIH: Monoclonal Anti-*Toxoplasma gondii* Surface Antigen 1 Protein, Clone T4 1E5 (produced *in vitro*), NR-50255), rabbit anti-AMA1 (1:2000, provided by the NIH Biodefense Proteomics Research Centers, NIAID, for distribution by BEI Resources, NIAID, NIH: AMA1, EBPC-3, Rabbit Polyclonal Antiserum, 3 Bleeds, Rabbit A3230, NR-4147); rabbit anti-ROP18 (1:5000, made in our laboratory). Detection was carried out using alkaline phosphatase-conjugated goat anti-rabbit secondary antibodies (Sigma-Aldrich), followed by development with the NBT/BCIP system (Promega Corporation, Fitchburg, WI, USA).

### 4.6. RT-PCR

Tachyzoites from RH and RHΔH2A.X strains were collected and stored in TRIzol reagent (ThermoFisher Scientific) at –80 °C until further use. Total RNA was extracted following the manufacturer’s instructions. Complementary DNA (cDNA) was synthesized using M-MLV reverse transcriptase (Promega Corporation) and oligo(dT) primers, according to the protocol provided with the enzyme. RT-PCR was then performed using specific primers to amplify H2A.X (Fw: 5’-AAAGGTGCAGGAGGTCGTAA-3’; Rv: 5’-CCTCCAAGGAACTTGCTCAG-3’) and tubulin (Fw: 5’-GAGCTCTTCTGCCTGGAA-3’; Rv: 5’-ATTTGGAGCCCACCGTCG-3’) transcripts.

### 4.7. Lytic cycle assays

For plaque assays, cells were grown in 6-well plates and infected with 200 RH, RHΔH2A.X or RHΔH2A.X-c-Myc-H2A.X complemented tachyzoites. After a 10-minute incubation on ice to promote parasite attachment, the plates were transferred to 37 °C for 1 hour to allow invasion. Subsequently, wells were washed with PBS and fresh medium was added. The plates were incubated at 37 °C for 7 days to allow plaque formation. After incubation, cells were fixed with cold 70% ethanol for 10 minutes at –20 °C and stained with 0.5% Crystal Violet for 5 minutes at room temperature. Excess dye was discarded, and plaques were imaged using a Discovery V2.0 SteREO Zeiss microscope at 422.0x magnification.

To assess invasion rates, cells were grown on slides and infected with exactly one parasite per cell. After 30 minutes of incubation, the monolayers were exhaustively washed and incubated for 18 h to allow replication, then fixed with cold 100% methanol for 7 minutes at -20 °C. IFA was then performed as described above. The number of vacuoles was counted in 50 randomly selected fields, and the number of vacuoles per field was plotted.

Gliding assays were performed by seeding extracellular parasites RH, RHΔH2A.X or RHΔH2A.X-c-Myc-H2A.X complemented onto poly-L-lysine-coated slides and incubated for 30 minutes at 37 °C. After incubation, parasites were fixed and processed for IFA using anti-SAG1 antibodies. The number of parasites exhibiting visible trails was quantified. The length of trails was measured using Image J.

Egress assays were performed as follows: cells grown on coverslips were infected with 10⁵ parasites and incubated for 26 to 30 hours. After incubation, monolayers were washed with serum-free DMEM and treated with 2 µM calcium ionophore (A23187, Sigma-Aldrich) in serum-free DMEM for 3 minutes to induce parasite egress. Following treatment, the ionophore was removed, and cells were fixed with cold 100% methanol for 7 minutes at -20 °C. Subsequently, IFA was performed according to the previously described protocol. Lysed and non-lysed vacuoles were quantified in 50 randomly selected fields.

Replication rates were obtained by IFA: hTERT cells were cultured on slides placed in 24-well plates and infected with 1×10⁵ RH or RHΔH2A.X tachyzoites (corresponding to a ratio of 1 parasite per cell). After a 1-hour incubation to allow invasion, the monolayers were washed with PBS and incubated for 20 hours. When evaluating the replication rate in the presence of genotoxic drugs, the monolayers were washed with PBS and incubated for 20 hours with DMEM supplemented with 0.1% dimethyl sulfoxide (DMSO, HCPT control), DMEM (MMS or HU controls), or the genotoxic agents HCPT (12 µM), methyl methanesulfonate (MMS, 50 µM) or hydroxyurea (HU 0.2 mM). Cells were fixed with 100% methanol for 7 minutes at 4 °C, and an IFA was performed to evaluate parasite replication by counting the number of parasites per vacuole in 100 randomly selected fields.

### 4.8. Flow cytometry and cell cycle analysis

hTERT cells were cultured to confluence in 6-well plates and subsequently infected with 10⁶ RH, RHΔH2A.X or RHΔH2A.X-c-Myc-H2A.X complemented tachyzoites per well. Once a significant number of mature parasitophorous vacuoles were observed, the infected monolayers were scrapped, and the cells were mechanically lysed by syringe forcing through needles of decreasing gauge (23G, 25G, 27G). Parasites were then purified by filtration through a 3 µm membrane, centrifuged at 2000 RPM for 10 minutes, washed with PBS, and fixed in 70% ethanol for 24 hours at -20 °C. After fixation, samples were centrifuged and washed with PBS containing 2% FBS. Following another centrifugation step, parasites were resuspended in 1 ml of PBS supplemented with RNase A (180 µg/ml, Roche, Manheim, Germany) and incubated at 37 °C for 10 minutes. Subsequently, propidium iodide (0.5 mg/ml, Sigma-Aldrich) was added, and samples were incubated for 10 minutes prior to acquisition on a BD FACS Calibur flow cytometer. Data was analyzed using FlowJo v10 software.

### 4.9. Ultrastructure Expansion Microscopy (U-ExM)

Ultrastructure expansion microscopy (U-ExM) was performed according to a previously described protocol [57], with minor modifications. Briefly, coverslips with infected cells were incubated for 5 hours at 37°C in anchoring solution (0.7 % acrylamide and 1 % formaldehyde in PBS), for covalent union of biomolecules to hydrogel. Polymerization started by introducing monomer solution (19% sodium acrylate, 10% ascorbic acid and 0.1% N,N′-metylenbisacrylamide in PBS), and incubated for 1 h at 37°C. After jellification, protein denaturation followed by incubation at 95°C for 1.5 h. After incubation, the gel was immersed in deionized water for 12-16 h for complete expansion. For staining, primary and secondary antibodies used were: anti centrin-1 (1:500), anti acetylated tubulin (1:500), anti TgEB1 (1:1500) and anti H2B.Z (1:250).

### 4.10. RNA-seq and transcriptomic analysis

At least 2 x10^8^ intracellular tachyzoites of RH and RHΔH2A.X were collected by syringe passage of scrapped cells, filtered and washed with PBS at 4°C. Four biological replicas of each were resuspended in RNA later (Sigma-Aldrich) and stored at -80°C until shipping to the HaploX facility (Hong Kong). RNA extraction was performed by HaploX service. Poly A libraries were prepared with samples between 1.0μg≤m<2.0μg, RIN≥6.5, 28S : 18S≥1.0 (concentration determined with Qubit Fluorometer and purity identification method: UNano-1000/1000 Spectrophotometer). PE150 sequencing (2x150, 20M reads/sample. 6G data) was obtained with Novaseq Xplus.

### 4.11. RNA-seq data processing and quantification

Raw paired-end Illumina reads were processed with the nf-core/rnaseq pipeline (v3.14.0) [58] implemented in Nextflow (v25.10.2) [59], executed with the Docker execution profile.

Adapter and quality trimming were performed with Trim Galore (v0.6.7) [60], based on Cutadapt (v3.4) [61], and read quality was assessed with FastQC (v0.12.1) [62] before and after trimming. Ribosomal RNA reads were depleted with SortMeRNA (v4.3.4) [63] using its default rRNA reference database. Genome alignment was skipped; gene expression was instead quantified by pseudo-alignment with Salmon (v1.10.1) [64] against an index built from the *Toxoplasma gondii* ME49 reference genome and annotation (ToxoDB release 68) [65]. Transcript-level abundances were summarized to the gene level with tximport[66]. Sample-level quality control (principal component analysis variance-stabilized counts) was performed within the pipeline with DESeq2 [67], and an aggregate QC report was generated with MultiQC [68]. The experiment comprised two conditions - the parental strain (RH, control), and H2A.X knockout (RH**Δ**H2A.X) - with four biological replicates each (n = 8 libraries). Two samples were excluded prior to differential expression analysis based on PCA outliers, KOH2Ax_3 and parRH_4 (the latter also showing the lowest mapping rate in its group, 64.9%). The final dataset comprised six samples: parRH_1–3, and KOH2Ax_1, 2 and 4 (Fig. S4).

### 4.12. Differential expression analysis

Length-scaled gene-level counts were analyzed in R (v4.5.3) with DESeq2 [67]. Genes were pre-filtered to those with ≥10 counts in ≥3 samples (7,224 of 8,618 genes retained). The RH**Δ**H2A.X-versus-parRH contrast was tested with the DESeq2 Wald test, with Benjamini-Hochberg (BH) correction. Genes with adjusted padj < 0.05 were considered significant for the broad screen used as input for functional enrichment; a stricter set additionally required |log₂FoldChange| ≥ 1. Gene identifiers were annotated with gene symbols and functional descriptions parsed from the ToxoDB-68 GFF file, and results were visualized as volcano plot with ggplot2 [69]. As an orthogonal, conservative filter, the same contrast (padj < 0.05 and |log₂FoldChange| ≥ 1) was additionally tested with edgeR (quasi-likelihood F-test) [70, 71] and limma-voom (empirical Bayes moderation) [72, 73], each under its standard null hypothesis (log₂FoldChange = 0) and the same design formula. For each direction, genes called significant by all three methods (DESeq2, edgeR, limma) were retained as a high-confidence intersection set. For downstream analysis significantly genes obtained from DESeq2 were used.

### 4.13. Functional enrichment analysis

Gene Ontology (GO) over-representation analysis (ORA) was performed separately on up- and down-regulated gene sets (broad DESeq2 screen, padj < 0.05) against the Biological Process, Molecular Function and Cellular Component ontologies via the ToxoDB REST API [65] (organism *T. gondii* ME49; evidence codes Computed and Curated; BH p < 0.05). GO Slim ORA was performed analogously against the ToxoDB Generic GO Slim gene set (goSubset = Yes). Redundancy among enriched GO terms was reduced with GOSemSim (Wang semantic similarity) [74], greedily retaining terms ranked by BH p-value at a pairwise similarity cutoff of 0.5, the default used by REVIGO [75]. KEGG pathway ORA was performed with clusterProfiler [76] against the *T. gondii* KEGG organism code tgo (BH p < 0.05).

### 4.14. Cell-cycle phase mapping of differentially expressed genes

Up- and down-regulated gene sets from the broad DESeq2 screen (RHΔH2A.X vs. RH; padj < 0.05, no log2FC threshold; n = 1,690 up, n = 1,682 down) were compared against the hourly cell-cycle transcriptome of thymidine-synchronized RH tachyzoites [41] (GEO GSE19092; ToxoDB dataset DS_c1a3dbb014), obtained programmatically via the ToxoDB/VEuPathDB WDK REST API (*GenesByMicroarray…CellCycle_RSRC* search; 13 hourly time points, 0–12 h). For each gene, the hour of maximal expression (RMA-normalized signal) was determined and compared against five overlapping phase windows: S(1) 0–3 h, M-C(1) 3–5 h, G1 4–8 h, S(2) 8–11 h, and M-C(2) 10–12 h (boundaries inclusive; genes peaking at shared boundaries were assigned to both adjacent phases). Genes lacking a valid microarray measurement (7.3% of up-regulated and 12.8% of down-regulated genes) were excluded from phase mapping. For each phase, the corresponding gene set was intersected separately with the up- and down-regulated DEG lists and visualized as a grouped bar plot (R v4.5.3, ggplot2).

### 4.15. Integration with H2A.X ChIP-seq data

For the integration we used H2A.X-Myc chromatin immunoprecipitation sequencing (ChIP-seq) peaks from Nardelli et al. [27] (GEO accession GSE87834). Peak-to-gene assignment was performed with ChIPseeker (v1.44.0) against the ToxoDB release 26 gene annotation, using a transcription start site (TSS)-proximal window of −1 to +200 bp to define promoter-associated peaks; gene identifiers (TGME49_XXXXXX) were used directly without liftover, as ToxoDB locus tags are stable across genome releases. All 15,618 called peaks in the source GFF were additionally classified by genomic context based on sequence identifier - main chromosome (TGME49_chr*), centromeric scaffold (ABPA02*), or subtelomeric scaffold (tgme49_asmbl*/TGG_*) - and their distance to the nearest annotated TSS was computed. Gene-level integration was restricted to the centromeric/subtelomeric peak-to-gene assignments, consistent with the reported enrichment of H2A.X at *Toxoplasma gondii* centromeres and telomeres [27], yielding 974 H2A.X-associated genes, of which 811 were represented among the 7,224 genes retained in the DESeq2 analysis after low-count filtering. H2A.X-bound genes were intersected with the RHΔH2Ax vs. RH differentially expressed gene (DEG) sets (DESeq2, padj < 0.05, either direction), and enrichment of H2A.X occupancy among up- and down-regulated genes was assessed independently by two-sided Fisher’s exact tests against the 7,224-gene background. Results were visualized as a three-way Venn diagram (H2A.X-bound genes vs. up- and down-regulated DEGs). For genome-track visualization, a bigWig file was generated from main-chromosome peaks using their reported peak scores. TSS reference points for deeptools were extracted from the release 68 GFF and stratified by gene category (H2A.X-bound, up-regulated, down-regulated, and transcriptionally stable). H2A.X signal enrichment around gene TSSs was profiled and visualized as metagene profile plots and heatmaps using deepTools (v3.5.4) computeMatrix, plotProfile, and plotHeatmap (reference-point mode, TSS ± 3 kb).

### 4.16. H2B.Z/H3K4me3/H2A.Z ChIP-seq and subtelomeric-location integration with RNA-seq

H2B.Z-myc, H3K4me3, and H2A.Z chromatin immunoprecipitation sequencing (ChIP-seq) peaks from Nardelli *et al. [27]* (GEO accession GSE104347) were integrated with the RHΔH2A.X vs. RH differential expression results. MACS2 broadPeak calls for H2B.Z (3 biological replicates, GSM2795870-72) and H2A.Z (2 biological replicates, GSM2795868-69), and narrowPeak calls for H3K4me3 (2 biological replicates, GSM2795873-74), were downloaded from GEO. Peak coordinates were confirmed to match the ToxoDB release 26 gene annotation coordinate system; gene identifiers (TGME49_XXXXXX) were used directly without liftover. Peak-to-gene assignment used a TSS-proximal promoter window of −1 to +200 bp (same convention as the H2A.X ChIP-seq analysis), built from the ToxoDB r26 GFF. A gene was considered bound by a given mark if its promoter window overlapped a called peak in a majority of that mark’s replicates (H2B.Z: ≥2 of 3; H3K4me3 and H2A.Z: 2 of 2), yielding 7,047 H2B.Z-bound genes (5,887 within the 7,224-gene DESeq2 background), 5,191 H3K4me3-bound genes (4,825), and 6,623 H2A.Z-bound genes (5,629). Unlike H2A.X, all three marks were assigned genome-wide (main chromosomes and centromeric/subtelomeric scaffolds), consistent with their expected promoter-proximal distribution. Genes were independently classified as subtelomeric if annotated on a subtelomeric scaffold (tgme49_asmbl*/TGG_* sequence identifiers) in the ToxoDB release 26 gene annotation, following the same scaffold-identity criterion used for H2A.X centromeric/subtelomeric classification but restricted to subtelomeric scaffolds (centromeric ABPA02* scaffolds excluded). This yielded 600 subtelomeric genes, of which 51 passed the DESeq2 expression filter.

For each of the four variables (H2B.Z occupancy, H3K4me3 occupancy, H2A.Z occupancy, and subtelomeric location), enrichment among upregulated and downregulated RHΔH2A.X vs. RH DEGs (DESeq2, *padj* < 0.05) was assessed by two-sided Fisher’s exact test against the 7,224-gene DESeq2 background, with Benjamini-Hochberg correction applied across the eight resulting tests. Results were visualized as volcano plots of the DESeq2 results with variable-positive genes highlighted by differential expression status (Fig.S4).

### 4.17. Gene set enrichment analysis (GSEA)

Genes were pre-ranked by log2 fold change x -log10(p-value) (DESeq2 results for RH**Δ**H2A.X-versus-RH, padj < 0.25 filter) and tested against GO Cellular Component (CC) gene sets (ToxoDB annotations, Computed + Curated evidence; min/max gene-set size 10-500) using clusterProfiler::GSEA [76] with Benjamini-Hochberg correction, alongside the same procedure for the Biological Process and Molecular Function ontologies. To obtain non-redundant summary of ontologies, term selection was automated in two steps. First, terms were excluded from the candidate pool if their leading edge was driven by a single gene or if the gene set was oversized. Second, Wang semantic similarity was performed (GOSemSim, cutoff = 0.5).

### 4.18. Statistics

Data were expressed as mean ± SD from three to four independent experiments. Data variations were analyzed with GraphPad Prism (8.01) software, using Student’s *t* test (*p ≤ 0.05, **p ≤ 0.01, and ***p ≤ 0.001).

For RNAseq, all statistical analyses were performed in R (v4.5.3) using DESeq2 [67], edgeR [70], limma [72], clusterProfiler [76], GOSemSim [73] and ggplot2 [69]. Unless otherwise stated, multiple-testing correction used the Benjamini-Hochberg method, and p/padj < 0.05 was considered statistically significant.

## Acknowledgements

1. C. Cristaldi (Fellow), A. Ganuza (Technician), A. A. Aparicio Arias (Fellow), A. M. Alonso (Researcher) S.O. Angel (Researcher) and L. Vanagas (Researcher) are members of CONICET. L. Gualano (Posdoc fellow) is funded by ANPCyT (Préstamo BID-PICT 2021-0169 to L.V.). S.O. Angel (Full), L. Gualano (JTP), A.M. Alonso (JTP), A. Ganuza (Assistant) and L. Vanagas (Adjunct) are Professors at Universidad Nacional General de San Martin (UNSAM). This work was funded by ANPCyT (Préstamo BID-PICT 2021-0169), UNSAM (80020250100027SM), NIH (R01 AI183372). The authors would like to thank Dr. Mariela S. Ferreira-Gomes (IQUIFIB, UBA-CONICET) for the kind gift of A23187 calcium ionophore.

## Data availability

The data that support the findings of this study are available from the corresponding author upon reasonable request. All RNA-seq raw sequencing data generated in this study have been deposited in the NCBI Gene Expression Omnibus (GEO) repository and will be made publicly available upon publication. Accession numbers will be provided prior to final acceptance. All other relevant data supporting the findings of this study are included within the manuscript and its Supporting Information files.

## Author Contributions

CC: Conceptualization, Investigation, Formal Analysis, Methodology, Writing – original draft, Writing – review & editing; SH: Conceptualization, Methodology, Investigation, Formal Analysis, Writing – review & editing; LG: Methodology, Investigation, Formal Analysis, Validation, Writing – review & editing; AG: Methodology, Investigation, Validation; AAAA: Methodology, Investigation, Validation; LDB: Methodology, Resources, Formal Analysis, Writing – review & editing; WJS: Conceptualization, Resources, Funding acquisition, Writing – review & editing; AMM: Methodology, Software, Formal Analysis, Data curation, Visualization, Writing – review & editing; SOA: Conceptualization, Resources, Supervision, Funding acquisition, Writing – review & editing; MEF: Conceptualization, Resources, Supervision, Writing – review & editing; LV: Conceptualization, Methodology, Investigation, Formal Analysis, Data curation, Supervision, Project administration, Writing supervision - original draft, Writing – review & editing.

## Supplementary material

**Supplementary Figure S1. Knockout generation strategy and validation of RHΔH2A.X clones.** (A) Gene editing strategy to generate the RHΔH2A.X line through the CRISPR-Cas9 technique. A guide RNA (gRNA) targeting the promoter region of the H2A.X gene was designed to direct Cas9 to the cleavage site. A replacement cassette containing the pyrimethamine resistance gene (DHFR, Dihydrofolate Reductase), flanked by homology arms located at the 5′ end, near the Cas9 cleavage site, and at the 3′ end of the H2A.X gene, was co-transfected. Homologous recombination-mediated repair led to insertion of the selection cassette in place of H2A.X. Recombinant parasites were selected with pyrimethamine and subsequently cloned. (B) Western blot analysis of RHΔH2A.X clones and RH strain. H2A.X and SAG1 (charge control) antibodies were used. A ∼17 kDa band corresponding to H2A.X was detected only in RH strain. (C) PCR on genomic DNA from clones 5 and 17. The absence of the H2A.X band (∼300 bp) and the presence of the DHFR band (∼3500 bp) indicate correct integration of the selection cassette and deletion of the H2A.X gene. Genomic DNA from RH parasites was used as a negative control, showing amplification only of H2A.X. (D) RT-PCR performed on cDNA from clones 5 and 17. Amplification of the expression control (tubulin) is observed in all samples, while H2A.X is not detected, confirming the absence of transcription. Genomic DNA and cDNA from RH parasites were included as positive controls, showing H2A.X amplification.

**Supplementary Figure S2. Analysis of cell division markers.** Representative IFA of parental RH and RHΔH2A.X tachyzoites stained with acetylated tubulin (white), Cpn60 (yellow) and DAPI (purple). Quantification showed no significant differences in the Cpn60 foci number of RHΔH2A.X compared to the parental.

**Supplementary Figure S3. Chromatin context of differentially expressed genes in RHΔH2A.X.** Volcano plots of RHΔH2A.X vs. RH transcriptomic comparison (log^2^ fold change vs. −log^10^ adjusted *p*-value). Genes whose promoters carry ChIP-seq peaks for H2B.Z (A), H2A.Z (B), or H3K4me3 (C) in the parental *T. gondii* strain [27] (GEO GSE104347) are highlighted: triangles indicate significantly differentially expressed genes (dark red: upregulated; dark blue: downregulated; *padj* < 0.05), orange circles indicate genes carrying the mark but not differentially expressed. Promoter occupancy was defined as ChIP-seq signal within a −1/+200 bp window around the TSS, present in the majority of available replicates (see Methods). (D) As in (A–C), highlighting the 51 subtelomeric genes (annotated on subtelomeric scaffolds in ToxoDB r26) that passed DESeq2 expression thresholds (out of 600 total). Subtelomeric analysis should be interpreted with caution given the limited background size (n = 51 genes).

**Supplementary Figure S4.** A Principal Component Analysis reveals clear differences at the transcriptomic level between the analyzed conditions. Three biological replicates for condition are presented. parRH: parental strain RH, KOH2Ax: RHΔH2A.X.

**Supplementary Table S1. Differential expression and GO Slim functional enrichment results for RHΔH2A.X versus RH**. The workbook comprises 18 sheets. **DESeq2_all**: full test results for all 7,224 expressed genes tested; **DESeq2_sig:** significantly detected genes (padj > 0.05). Up and Down regulated genes (|LFC| > 0 | |LFC|< 0) were highlighted as “UP” or “DOWN” at column up/down; **DESeq2_lfc1_sig**: 461 genes significant by DESeq2 (padj < 0.05, |LFC| ≥ 1); **edgeR_all**: full edgeR glmQLFTest results for all 7,224 genes; **edgeR_sig**: 494 genes significant by edgeR (FDR < 0.05, |LFC| ≥ 1); **limma_all**: full limma-voom results for all 7,224 genes; **limma_sig**: 443 genes significant by limma (adj.P < 0.05, |log2FC| ≥ 1); **Intersection_up**: 52 up-regulated genes called significant by all three methods (DESeq2, edgeR, limma); **Intersection_dn**: 329 down-regulated genes called significant by all three methods; **GOslim_BP_up** / **GOslim_BP_down**: GO Slim Biological Process terms enriched among up- (16 terms) and down-regulated genes (6 terms); **GOslim_MF_up** / **GOslim_MF_down**: GO Slim Molecular Function terms enriched among up- (12 terms) and down-regulated genes (4 terms); **GOslim_CC_up** / **GOslim_CC_down**: GO Slim Cellular Component terms enriched among up- (14 terms) and down-regulated genes (3 terms); **GSEA_go_cc_simplified**: the five robust, non-redundant GO Cellular Component terms from pre-ranked GSEA; **CHIP_direct_targets_up**: 171 H2A.X ChIP-seq target genes that are also significantly up-regulated; **CHIP_direct_targets_down**: 201 H2A.X ChIP-seq target genes that are also significantly down-regulated.

**Supplementary Table S2. Differentially expressed genes involved in *Toxoplasma gondii* invasion, gliding, and adhesion.** Overview and full dataset of differentially expressed genes grouped by functional categories (micronemes, rhoptries, dense granules, surface antigens, proteases, and signaling/transport components). The file includes a category summary tab ("Summary") and the detailed gene catalog ("Gene_List") containing ToxoDB gene IDs, gene symbols, descriptions, log_2_ FoldChange , adjusted *p*-values (*padj*), and expression directionality (UP/DOWN).

**Supplementary Table S3. Differentially expressed genes related to DNA replication, mitotic machinery, and protein synthesis.** Overview and full dataset of differentially expressed genes associated with cell cycle progression, genome replication, chromosome segregation, and translational machinery. The file includes a category summary tab ("Summary") and the detailed gene catalog ("Gene_List") containing ToxoDB gene IDs, gene symbols, descriptions, log_2_ FoldChange, adjusted *p*-values (*padj*), and expression directionality (UP/DOWN) across functional categories (DNA replication factors, kinetochore, condensin, centrosome/centriole components, Aurora-type kinases, and ribosomal proteins).

## References

1. Guo M, Dubey JP, Hill D, Buchanan RL, Gamble HR, Jones JL, et al. Prevalence and risk factors for Toxoplasma gondii infection in meat animals and meat products destined for human consumption. J Food Prot. 2015;78(2):457–76. Epub 2015/02/25. doi: 10.4315/0362-028X.JFP-14-328. PubMed PMID: 25710166.

2. Basavaraju A. Toxoplasmosis in HIV infection: An overview. Trop Parasitol. 2016;6(2):129–35. Epub 2016/10/11. doi: 10.4103/2229-5070.190817. PubMed PMID: 27722101; PubMed Central PMCID: PMCPMC5048699.

3. Tenter AM, Heckeroth AR, Weiss LM. Toxoplasma gondii: from animals to humans. Int J Parasitol. 2000;30(12-13):1217–58. PubMed PMID: 11113252; PubMed Central PMCID: PMCPMC3109627.

4. Angel SO, Vanagas L, Alonso AM. Mechanisms of adaptation and evolution in Toxoplasma gondii. Molecular and biochemical parasitology. 2024;258:111615. Epub 2024/02/15. doi: 10.1016/j.molbiopara.2024.111615. PubMed PMID: 38354788.

5. Waldman BS, Schwarz D, Wadsworth MH, 2nd, Saeij JP, Shalek AK, Lourido S. Identification of a Master Regulator of Differentiation in Toxoplasma. Cell. 2020;180(2):359–72 e16. doi: 10.1016/j.cell.2019.12.013. PubMed PMID: 31955846; PubMed Central PMCID: PMC6978799.

6. Licon MH, Giuliano CJ, Chan AW, Chakladar S, Eberhard JN, Shallberg LA, et al. Author Correction: A positive feedback loop controls Toxoplasma chronic differentiation. Nat Microbiol. 2023. Epub 2023/08/16. doi: 10.1038/s41564-023-01467-y. PubMed PMID: 37582867.

7. Sokol-Borrelli SL, Reilly SM, Holmes MJ, Orchanian SB, Massmann MD, Sharp KG, et al. A transcriptional network required for bradyzoite development in Toxoplasma gondii is dispensable for recrudescent disease. Nature communications. 2023;14(1):6078. Epub 2023/09/29. doi: 10.1038/s41467-023-40948-w. PubMed PMID: 37770433; PubMed Central PMCID: PMCPMC10539341.

8. Luger K, Mader AW, Richmond RK, Sargent DF, Richmond TJ. Crystal structure of the nucleosome core particle at 2.8 A resolution. Nature. 1997;389(6648):251–60. doi: 10.1038/38444. PubMed PMID: 9305837.

9. Henikoff S, Furuyama T, Ahmad K. Histone variants, nucleosome assembly and epigenetic inheritance. Trends Genet. 2004;20(7):320–6. Epub 2004/06/29. doi: 10.1016/j.tig.2004.05.004. PubMed PMID: 15219397.

10. Jin J, Cai Y, Li B, Conaway RC, Workman JL, Conaway JW, et al. In and out: histone variant exchange in chromatin. Trends in biochemical sciences. 2005;30(12):680–7. Epub 2005/11/01. doi: 10.1016/j.tibs.2005.10.003. PubMed PMID: 16257529.

11. Guardamagna I, Lonati L, Iaria O, Mentana A, Parodi D, Peterlin G, et al. Temporal and spatial patterns of gammaH2AX signaling in different human cells after exposure to X-rays and UV-C light. Radiat Environ Biophys. 2026;65(2):781–91. Epub 2026/05/22. doi: 10.1007/s00411-026-01220-z. PubMed PMID: 42171732; PubMed Central PMCID: PMCPMC13303546.

12. de Feraudy S, Revet I, Bezrookove V, Feeney L, Cleaver JE. A minority of foci or pan-nuclear apoptotic staining of gammaH2AX in the S phase after UV damage contain DNA double-strand breaks. Proc Natl Acad Sci U S A. 2010;107(15):6870–5. Epub 2010/03/31. doi: 10.1073/pnas.1002175107. PubMed PMID: 20351298; PubMed Central PMCID: PMCPMC2872460.

13. Dhuppar S, Roy S, Mazumder A. gammaH2AX in the S Phase after UV Irradiation Corresponds to DNA Replication and Does Not Report on the Extent of DNA Damage. Mol Cell Biol. 2020;40(20). Epub 2020/08/12. doi: 10.1128/MCB.00328-20. PubMed PMID: 32778572; PubMed Central PMCID: PMCPMC7523655.

14. Turinetto V, Giachino C. Multiple facets of histone variant H2AX: a DNA double-strand-break marker with several biological functions. Nucleic Acids Res. 2015;43(5):2489–98. Epub 2015/02/26. doi: 10.1093/nar/gkv061. PubMed PMID: 25712102; PubMed Central PMCID: PMCPMC4357700.

15. Shimada M, Goshima T, Matsuo H, Johmura Y, Haruta M, Murata K, et al. Essential role of autoactivation circuitry on Aurora B-mediated H2AX-pS121 in mitosis. Nature communications. 2016;7:12059. Epub 2016/07/09. doi: 10.1038/ncomms12059. PubMed PMID: 27389782; PubMed Central PMCID: PMCPMC4941122.

16. Rogakou EP, Pilch DR, Orr AH, Ivanova VS, Bonner WM. DNA double-stranded breaks induce histone H2AX phosphorylation on serine 139. J Biol Chem. 1998;273(10):5858–68. PubMed PMID: 9488723.

17. Redon C, Pilch D, Rogakou E, Sedelnikova O, Newrock K, Bonner W. Histone H2A variants H2AX and H2AZ. Current opinion in genetics & development. 2002;12(2):162–9. PubMed PMID: 11893489.

18. Sullivan WJ, Jr., Naguleswaran A, Angel SO. Histones and histone modifications in protozoan parasites. Cellular microbiology. 2006;8(12):1850–61. Epub 2006/10/10. doi: 10.1111/j.1462-5822.2006.00818.x. PubMed PMID: 17026479.

19. Goyal M, Heinberg A, Mitesser V, Kandelis-Shalev S, Singh BK, Dzikowski R. Phosphorylation of the Canonical Histone H2A Marks Foci of Damaged DNA in Malaria Parasites. mSphere. 2021;6(1). Epub 2021/01/15. doi: 10.1128/mSphere.01131-20. PubMed PMID: 33441412; PubMed Central PMCID: PMCPMC7845613.

20. McLaughlin E, Zavala Martinez MG, Dujeancourt-Henry A, Chaze T, Gianetto QG, Matondo M, et al. Phosphoproteomic analysis of the response to DNA damage in Trypanosoma brucei. J Biol Chem. 2024;300(9):107657. Epub 2024/08/12. doi: 10.1016/j.jbc.2024.107657. PubMed PMID: 39128729; PubMed Central PMCID: PMCPMC11408851.

21. Glover L, Horn D. Trypanosomal histone gammaH2A and the DNA damage response. Molecular and biochemical parasitology. 2012;183(1):78–83. Epub 2012/02/23. doi: 10.1016/j.molbiopara.2012.01.008. PubMed PMID: 22353557; PubMed Central PMCID: PMCPMC3334830.

22. Dalmasso MC, Onyango DO, Naguleswaran A, Sullivan WJ, Jr., Angel SO. Toxoplasma H2A variants reveal novel insights into nucleosome composition and functions for this histone family. J Mol Biol. 2009;392(1):33–47. doi: 10.1016/j.jmb.2009.07.017. PubMed PMID: 19607843; PubMed Central PMCID: PMCPMC2734877.

23. Munera Lopez J, Ganuza A, Bogado SS, Munoz D, Ruiz DM, Sullivan WJ, Jr., et al. Evaluation of ATM Kinase Inhibitor KU-55933 as Potential Anti-Toxoplasma gondii Agent. Frontiers in cellular and infection microbiology. 2019;9:26. doi: 10.3389/fcimb.2019.00026. PubMed PMID: 30815397; PubMed Central PMCID: PMC6381018.

24. Cristaldi C, Saldarriaga Cartagena AM, Ganuza A, Sullivan WJ, Jr., Angel SO, Vanagas L. Evaluation of topotecan and 10-hydroxycamptothecin on Toxoplasma gondii: Implications on baseline DNA damage and repair efficiency. Int J Parasitol Drugs Drug Resist. 2023;23:120–9. Epub 2023/12/04. doi: 10.1016/j.ijpddr.2023.11.004. PubMed PMID: 38043188; PubMed Central PMCID: PMCPMC10730954.

25. Angel SO, Vanagas L, Ruiz DM, Cristaldi C, Saldarriaga Cartagena AM, Sullivan WJJ. Emerging Therapeutic Targets Against Toxoplasma gondii: Update on DNA Repair Response Inhibitors and Genotoxic Drugs. Front Cell Infect Microbiol. 2020;10(289). doi: doi: 10.3389/fcimb.2020.00289.

26. Saldarriaga Cartagena AM, Aparicio Arias A, Cristaldi C, Ganuza A, Gonzalez MM, Corvi MM, et al. Toxoplasma gondii RAD51 recombinase is required to overcome DNA replication stress and its inactivation leads to bradyzoite differentiation. DNA Repair (Amst). 2025;152:103882. Epub 2025/08/11. doi: 10.1016/j.dnarep.2025.103882. PubMed PMID: 40784138.

27. Nardelli SC, Silmon de Monerri NC, Vanagas L, Wang X, Tampaki Z, Sullivan WJ, Jr., et al. Genome-wide localization of histone variants in Toxoplasma gondii implicates variant exchange in stage-specific gene expression. BMC genomics. 2022;23(1):128. Epub 2022/02/16. doi: 10.1186/s12864-022-08338-6. PubMed PMID: 35164683; PubMed Central PMCID: PMCPMC8842566.

28. Contreras SM, Zambrano Siri RT, Rivera EM, Cristaldi C, Kamenetzky L, Kim K, et al. Architecture, Chromatin and Gene Organization of Toxoplasma gondii Subtelomeres. Epigenomes. 2022;6(3). Epub 2022/09/23. doi: 10.3390/epigenomes6030029. PubMed PMID: 36135316; PubMed Central PMCID: PMCPMC9498087.

29. Sidik SM, Huet D, Ganesan SM, Huynh MH, Wang T, Nasamu AS, et al. A Genome-wide CRISPR Screen in Toxoplasma Identifies Essential Apicomplexan Genes. Cell. 2016;166(6):1423–35 e12. doi: 10.1016/j.cell.2016.08.019. PubMed PMID: 27594426; PubMed Central PMCID: PMC5017925.

30. Giuliano CJ, Wei KJ, Harling FM, Waldman BS, Farringer MA, Boydston EA, et al. CRISPR-based functional profiling of the Toxoplasma gondii genome during acute murine infection. Nat Microbiol. 2024;9(9):2323–43. Epub 2024/07/09. doi: 10.1038/s41564-024-01754-2. PubMed PMID: 38977907; PubMed Central PMCID: PMCPMC11811839.

31. Vanagas L, Munoz D, Cristaldi C, Ganuza A, Najera R, Bonardi MC, et al. Histone variant H2B.Z acetylation is necessary for maintenance of Toxoplasma gondii biological fitness. Biochim Biophys Acta Gene Regul Mech. 2023;1866(3):194943. Epub 2023/05/23. doi: 10.1016/j.bbagrm.2023.194943. PubMed PMID: 37217032.

32. Kuo LJ, Yang LX. Gamma-H2AX - a novel biomarker for DNA double-strand breaks. In Vivo. 2008;22(3):305–9. Epub 2008/07/10. PubMed PMID: 18610740.

33. Paull TT, Rogakou EP, Yamazaki V, Kirchgessner CU, Gellert M, Bonner WM. A critical role for histone H2AX in recruitment of repair factors to nuclear foci after DNA damage. Curr Biol. 2000;10(15):886–95. Epub 2000/08/26. doi: 10.1016/s0960-9822(00)00610-2. PubMed PMID: 10959836.

34. Hsiang YH, Lihou MG, Liu LF. Arrest of replication forks by drug-stabilized topoisomerase I-DNA cleavable complexes as a mechanism of cell killing by camptothecin. Cancer Res. 1989;49(18):5077–82. Epub 1989/09/15. PubMed PMID: 2548710.

35. Shaw AE, Mihelich MN, Whitted JE, Reitman HJ, Timmerman AJ, Tehseen M, et al. Revised mechanism of hydroxyurea-induced cell cycle arrest and an improved alternative. Proc Natl Acad Sci U S A. 2024;121(42):e2404470121. Epub 2024/10/07 22:18. doi: 10.1073/pnas.2404470121. PubMed PMID: 39374399; PubMed Central PMCID: PMCPMC11494364.

36. Sibley LD. Intracellular parasite invasion strategies. Science. 2004;304(5668):248–53. Epub 2004/04/10. doi: 10.1126/science.1094717. PubMed PMID: 15073368.

37. Francia ME, Striepen B. Cell division in apicomplexan parasites. Nat Rev Microbiol. 2014;12(2):125–36. Epub 2014/01/05. doi: 10.1038/nrmicro3184. PubMed PMID: 24384598.

38. Chen CT, Kelly M, Leon J, Nwagbara B, Ebbert P, Ferguson DJ, et al. Compartmentalized Toxoplasma EB1 bundles spindle microtubules to secure accurate chromosome segregation. Mol Biol Cell. 2015;26(25):4562–76. Epub 2015/10/16. doi: 10.1091/mbc.E15-06-0437. PubMed PMID: 26466679; PubMed Central PMCID: PMCPMC4678015.

39. Striepen B, Crawford MJ, Shaw MK, Tilney LG, Seeber F, Roos DS. The plastid of Toxoplasma gondii is divided by association with the centrosomes. J Cell Biol. 2000;151(7):1423–34. Epub 2001/01/03. doi: 10.1083/jcb.151.7.1423. PubMed PMID: 11134072; PubMed Central PMCID: PMCPMC2150670.

40. Blader IJ, Coleman BI, Chen CT, Gubbels MJ. Lytic Cycle of Toxoplasma gondii: 15 Years Later. Annual review of microbiology. 2015;69:463–85. Epub 2015/09/04. doi: 10.1146/annurev-micro-091014-104100. PubMed PMID: 26332089; PubMed Central PMCID: PMCPMC4659696.

41. Behnke MS, Wootton JC, Lehmann MM, Radke JB, Lucas O, Nawas J, et al. Coordinated progression through two subtranscriptomes underlies the tachyzoite cycle of Toxoplasma gondii. PloS one. 2010;5(8):e12354. doi: 10.1371/journal.pone.0012354. PubMed PMID: 20865045; PubMed Central PMCID: PMC2928733.

42. Subramanian A, Tamayo P, Mootha VK, Mukherjee S, Ebert BL, Gillette MA, et al. Gene set enrichment analysis: a knowledge-based approach for interpreting genome-wide expression profiles. Proc Natl Acad Sci U S A. 2005;102(43):15545–50. Epub 2005/10/04. doi: 10.1073/pnas.0506580102. PubMed PMID: 16199517; PubMed Central PMCID: PMCPMC1239896.

43. Eliezer Y, Argaman L, Kornowski M, Roniger M, Goldberg M. Interplay between the DNA damage proteins MDC1 and ATM in the regulation of the spindle assembly checkpoint. J Biol Chem. 2014;289(12):8182–93. Epub 2014/02/11. doi: 10.1074/jbc.M113.532739. PubMed PMID: 24509855; PubMed Central PMCID: PMCPMC3961647.

44. Musacchio A, Salmon ED. The spindle-assembly checkpoint in space and time. Nature reviews Molecular cell biology. 2007;8(5):379–93. Epub 2007/04/12. doi: 10.1038/nrm2163. PubMed PMID: 17426725.

45. Yuan J, Adamski R, Chen J. Focus on histone variant H2AX: to be or not to be. FEBS letters. 2010;584(17):3717–24. Epub 2010/05/25. doi: 10.1016/j.febslet.2010.05.021. PubMed PMID: 20493860; PubMed Central PMCID: PMCPMC3695482.

46. Fox BA, Ristuccia JG, Gigley JP, Bzik DJ. Efficient gene replacements in Toxoplasma gondii strains deficient for nonhomologous end joining. Eukaryot Cell. 2009;8(4):520–9. doi: 10.1128/EC.00357-08. PubMed PMID: 19218423; PubMed Central PMCID: PMCPMC2669201.

47. Pavlou G, Biesaga M, Touquet B, Lagal V, Balland M, Dufour A, et al. Toxoplasma Parasite Twisting Motion Mechanically Induces Host Cell Membrane Fission to Complete Invasion within a Protective Vacuole. Cell host & microbe. 2018;24(1):81–96 e5. Epub 2018/07/17. doi: 10.1016/j.chom.2018.06.003. PubMed PMID: 30008293.

48. Birnbaum J, Flemming S, Reichard N, Soares AB, Mesen-Ramirez P, Jonscher E, et al. A genetic system to study Plasmodium falciparum protein function. Nature methods. 2017;14(4):450–6. Epub 2017/03/14. doi: 10.1038/nmeth.4223. PubMed PMID: 28288121.

49. Bougdour A, Durandau E, Brenier-Pinchart MP, Ortet P, Barakat M, Kieffer S, et al. Host cell subversion by Toxoplasma GRA16, an exported dense granule protein that targets the host cell nucleus and alters gene expression. Cell host & microbe. 2013;13(4):489–500. Epub 2013/04/23. doi: 10.1016/j.chom.2013.03.002. PubMed PMID: 23601110.

50. Marq JB, Gosetto M, Altenried A, Vadas O, Maco B, ao Cytokinetic abscission in Toxoplasma gondii is governed by protein phosphatase 2A and the daughter cell scaffold complex. The EMBO journal. 2024;43(17):3752–86. Epub 2024/07/16. doi: 10.1038/s44318-024-00171-9. PubMed PMID: 39009675; PubMed Central PMCID: PMCPMC11377541.

51. Reese ML, Zeiner GM, Saeij JP, Boothroyd JC, Boyle JP. Polymorphic family of injected pseudokinases is paramount in Toxoplasma virulence. Proc Natl Acad Sci U S A. 2011;108(23):9625–30. Epub 2011/03/26. doi: 10.1073/pnas.1015980108. PubMed PMID: 21436047; PubMed Central PMCID: PMCPMC3111280.

52. Cortes A, Carret C, Kaneko O, Yim Lim BY, Ivens A, Holder AA. Epigenetic silencing of Plasmodium falciparum genes linked to erythrocyte invasion. PLoS pathogens. 2007;3(8):e107. Epub 2007/08/07. doi: 10.1371/journal.ppat.0030107. PubMed PMID: 17676953; PubMed Central PMCID: PMCPMC1937010.

53. Rayner JC, Vargas-Serrato E, Huber CS, Galinski MR, Barnwell JW. A Plasmodium falciparum homologue of Plasmodium vivax reticulocyte binding protein (PvRBP1) defines a trypsin-resistant erythrocyte invasion pathway. The Journal of experimental medicine. 2001;194(11):1571–81. Epub 2001/12/06. doi: 10.1084/jem.194.11.1571. PubMed PMID: 11733572; PubMed Central PMCID: PMCPMC2193530.

54. Farwell DG, Shera KA, Koop JI, Bonnet GA, Matthews CP, Reuther GW, et al. Genetic and epigenetic changes in human epithelial cells immortalized by telomerase. The American journal of pathology. 2000;156(5):1537–47. Epub 2000/05/04. doi: 10.1016/S0002-9440(10)65025-0. PubMed PMID: 10793065; PubMed Central PMCID: PMCPMC1876907.

55. Donald RG, Roos DS. Gene knock-outs and allelic replacements in Toxoplasma gondii: HXGPRT as a selectable marker for hit-and-run mutagenesis. Molecular and biochemical parasitology. 1998;91(2):295–305. Epub 1998/05/05. doi: 10.1016/s0166-6851(97)00210-7. PubMed PMID: 9566522.

56. Bogado SS, Dalmasso MC, Ganuza A, Kim K, Sullivan WJ, Jr., Angel SO, et al. Canonical histone H2Ba and H2A.X dimerize in an opposite genomic localization to H2A.Z/H2B.Z dimers in Toxoplasma gondii. Molecular and biochemical parasitology. 2014;197(1-2):36–42. doi: 10.1016/j.molbiopara.2014.09.009. PubMed PMID: 25286383; PubMed Central PMCID: PMC4254167.

57. Tosetti N, Dos Santos Pacheco N, Bertiaux E, Maco B, Bournonville L, Hamel V, et al. Essential function of the alveolin network in the subpellicular microtubules and conoid assembly in Toxoplasma gondii. eLife. 2020;9. Epub 2020/05/08. doi: 10.7554/eLife.56635. PubMed PMID: 32379047; PubMed Central PMCID: PMCPMC7228768.

58. Ewels PA, Peltzer A, Fillinger S, Patel H, Alneberg J, Wilm A, et al. The nf-core framework for community-curated bioinformatics pipelines. Nature biotechnology. 2020;38(3):276–8. Epub 2020/02/15. doi: 10.1038/s41587-020-0439-x. PubMed PMID: 32055031.

59. Di Tommaso P, Chatzou M, Floden EW, Barja PP, Palumbo E, Notredame C. Nextflow enables reproducible computational workflows. Nature biotechnology. 2017;35(4):316–9. Epub 2017/04/12. doi: 10.1038/nbt.3820. PubMed PMID: 28398311.

60. Krueger F. Trim Galore. 2019.

61. Martin M. Cutadapt removes adapter sequences from high-throughput sequencing reads EMBnetjournal. 2011;17. doi: 10.14806/ej.17.1.200

62. Andrews S. FASTQC. A quality control tool for high throughput sequence data. 2010.

63. Kopylova E, Noe L, Touzet H. SortMeRNA: fast and accurate filtering of ribosomal RNAs in metatranscriptomic data. Bioinformatics. 2012;28(24):3211–7. Epub 2012/10/17. doi: 10.1093/bioinformatics/bts611. PubMed PMID: 23071270.

64. Patro R, Duggal G, Love MI, Irizarry RA, Kingsford C. Salmon provides fast and bias-aware quantification of transcript expression. Nature methods. 2017;14(4):417–9. Epub 2017/03/07. doi: 10.1038/nmeth.4197. PubMed PMID: 28263959; PubMed Central PMCID: PMCPMC5600148.

65. Amos B, Aurrecoechea C, Barba M, Barreto A, Basenko EY, Bazant W, et al. VEuPathDB: the eukaryotic pathogen, vector and host bioinformatics resource center. Nucleic Acids Res. 2022;50(D1):D898–D911. Epub 2021/11/01. doi: 10.1093/nar/gkab929. PubMed PMID: 34718728; PubMed Central PMCID: PMCPMC8728164.

66. Soneson C, Love MI, Robinson MD. Differential analyses for RNA-seq: transcript-level estimates improve gene-level inferences. F1000Res. 2015;4:1521. Epub 2016/03/01. doi: 10.12688/f1000research.7563.2. PubMed PMID: 26925227; PubMed Central PMCID: PMCPMC4712774.

67. Love MI, Huber W, Anders S. Moderated estimation of fold change and dispersion for RNA-seq data with DESeq2. Genome biology. 2014;15(12):550. Epub 2014/12/18. doi: 10.1186/s13059-014-0550-8. PubMed PMID: 25516281; PubMed Central PMCID: PMCPMC4302049.

68. Ewels P, Magnusson M, Lundin S, Kaller M. MultiQC: summarize analysis results for multiple tools and samples in a single report. Bioinformatics. 2016;32(19):3047–8. Epub 2016/06/18. doi: 10.1093/bioinformatics/btw354. PubMed PMID: 27312411; PubMed Central PMCID: PMCPMC5039924.

69. Wickham H. ggplot2: Elegant Graphics for Data Analysis. Springer-Verlag. 2016. doi: 10.1007/978-3-319-24277-4_9

70. Robinson MD, McCarthy DJ, Smyth GK. edgeR: a Bioconductor package for differential expression analysis of digital gene expression data. Bioinformatics. 2010;26(1):139–40. Epub 2009/11/17. doi: 10.1093/bioinformatics/btp616. PubMed PMID: 19910308; PubMed Central PMCID: PMCPMC2796818.

71. Lun AT, Chen Y, Smyth GK. It’s DE-licious: A Recipe for Differential Expression Analyses of RNA-seq Experiments Using Quasi-Likelihood Methods in edgeR. Methods in molecular biology. 2016;1418:391–416. Epub 2016/03/24. doi: 10.1007/978-1-4939-3578-9_19. PubMed PMID: 27008025.

72. Ritchie ME, Phipson B, Wu D, Hu Y, Law CW, Shi W, et al. limma powers differential expression analyses for RNA-sequencing and microarray studies. Nucleic Acids Res. 2015;43(7):e47. Epub 2015/01/22. doi: 10.1093/nar/gkv007. PubMed PMID: 25605792; PubMed Central PMCID: PMCPMC4402510.

73. Law CW, Chen Y, Shi W, Smyth GK. voom: Precision weights unlock linear model analysis tools for RNA-seq read counts. Genome biology. 2014;15(2):R29. Epub 2014/02/04. doi: 10.1186/gb-2014-15-2-r29. PubMed PMID: 24485249; PubMed Central PMCID: PMCPMC4053721.

74. Yu G, Li F, Qin Y, Bo X, Wu Y, Wang S. GOSemSim: an R package for measuring semantic similarity among GO terms and gene products. Bioinformatics. 2010;26(7):976–8. Epub 2010/02/25. doi: 10.1093/bioinformatics/btq064. PubMed PMID: 20179076.

75. Supek F, Bosnjak M, Skunca N, Smuc T. REVIGO summarizes and visualizes long lists of gene ontology terms. PloS one. 2011;6(7):e21800. Epub 2011/07/27. doi: 10.1371/journal.pone.0021800. PubMed PMID: 21789182; PubMed Central PMCID: PMCPMC3138752.

76. Wu T, Hu E, Xu S, Chen M, Guo P, Dai Z, et al. clusterProfiler 4.0: A universal enrichment tool for interpreting omics data. Innovation (Camb). 2021;2(3):100141. Epub 2021/09/25. doi: 10.1016/j.xinn.2021.100141. PubMed PMID: 34557778; PubMed Central PMCID: PMCPMC8454663.

